# Evolutionary survival strategies of the female giant panda: optimizing energy resources and expenditure prior to pregnancy by postponing corpus luteum reactivation

**DOI:** 10.1101/2022.06.28.497916

**Authors:** Jella Wauters, Kirsten S. Wilson, Tim Bouts, Catherine Vancsok, Baptiste Mulot, Antoine Leclerc, Marko Haapakoski, José Kok, Ragnar Kühne, Andreas Ochs, Alan S. McNeilly, Mick Rae, Ruth Andrew, W. Colin Duncan, Simon J. Girling, Quang Zhou, Rengui Li, Yingmin Zhou, Kailai Cai, Yuliang Liu, Rong Hou, Iain Valentine, Thomas B. Hildebrandt, Desheng Li, Lynn Vanhaecke

## Abstract

The giant panda reproductive physiology shows important similarities with at least six of the eight existing bear species: the occurrence of diapause and/or pseudopregnancy is commonly described in bears. Nevertheless, significant differences including the earlier breeding season with - in general - a single estrus, a shorter delay of implantation and a more variable birth season, indicate an evolutionary adaptation from the seasonal reproductive traits described in hibernating bear species.

In this study we aimed to determine true pregnancy length for giant pandas and to open the discussion on the peculiarities of giant panda reproductive biology, more specifically focusing on the rationale behind their short-seasoned reproductive cycle compared to the other bear species.

For this purpose, metabolic (body weight and fecal output) profiles were matched with endocrine changes, mainly urinary progesterone metabolites, in 5 pregnant, 8 non-birth and 6 pseudopregnant cycles of 6 female giant pandas.

Pregnancy in giant pandas lasts only 42 days from early reactivation of the corpora lutea (CLs) until birth. In addition, our findings urged the need to redefine the generally accepted biphasic progesterone profile into a triphasic primary progesterone rise (corpus luteum dormancy (CLD) I, II and III) prior to entering the active luteal phase (= secondary progesterone rise). Two episodes of progesterone increase (CLDII: 81.20 ± 5.85 days versus CLDIII: 60.80 ± 3.83 days prior to birth for pregnant cycles) were identified, respectively corresponding to CL reactivation (74-88 days prior to birth) and implantation (± 60 days prior to birth) in other bear species. The progesterone concentration during CLDIII was higher in pregnant cycles, indicating a potential communication between maternal tissues and blastocyst(s) enhancing progesterone concentrations and thus allowing optimal priming of uterine tissues to better prepare for blastocyst reactivation/development. Compared with other bear species, giant pandas seem to shorten the active luteal phase, and thus pregnancy, by approximately 30 days by actively postponing CL reactivation.

Potential mechanisms in play overruling and suppressing the evolutionary conserved photoperiodical triggers of CL reactivation are discussed while a parallel study will further elaborate on the CL dynamics during giant panda gestation.

## Introduction

Giant panda reproductive biology largely resembles that of most other bear species, including diapause (e.g. American, Asiatic and Japanese black bear; polar bear; (Hokkaido) brown bear) and pseudopregnancy (e.g. American, Asiatic and Japanese black bear; polar bear; (Hokkaido) brown bear, sun bear). However, there are also significant differences: earlier and longer female breeding season in late winter/early spring versus late spring/summer, shorter delay of implantation of on average 2-3 months compared with up to 6 months in the other bear species and earlier and more variable birth season, i.e. summer versus winter, as compared to almost all obligate hibernating (e.g. black and brown) and facultative or non-hibernating (e.g. polar) bear species (Fig. 1).

**Fig 1:**
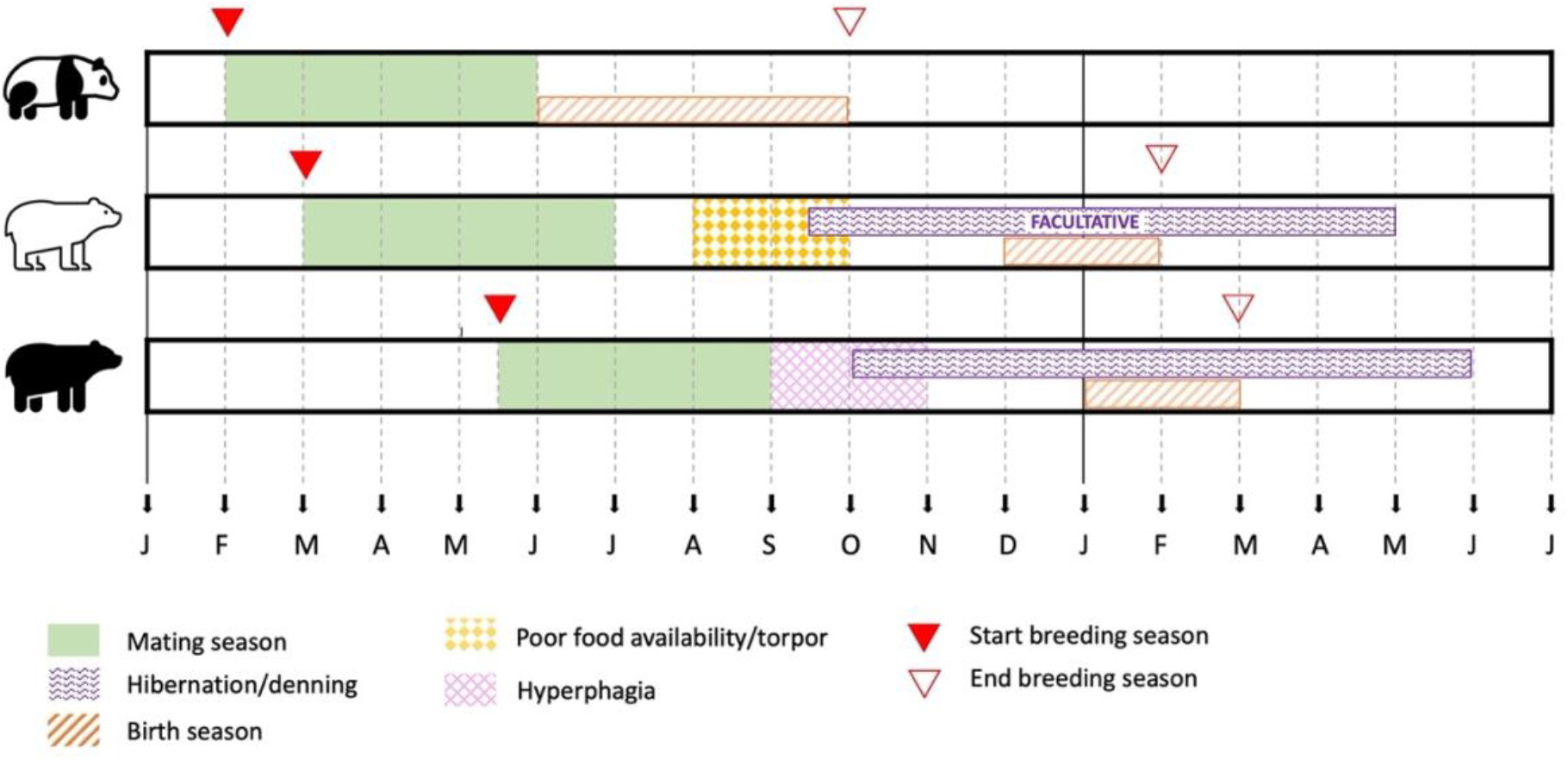
Generalized comparative overview of the reproductive cycle of bear species with key examples such as the giant panda, polar bear and American black bear. The breeding season of giant pandas is shorter (winter/spring - early autumn) compared to (facultative) hibernating species (spring/summer – winter), with an earlier onset of mating season, shorter diapause, absence of hibernation and more variable birth season. In polar bears, only (pseudo)pregnant female polar bears undergo hibernation. Black bears undergo a period of hyperphagia prior to the onset of hibernation, while female pregnant polar bears are forced into hibernation after a period of severe food scarcity typically occurring from late summer onwards.

While the giant panda is a monoestrous spontaneous ovulator, most of the other investigated bear species are polyoestrous (i.e. American black bear, polar bear, brown bear and sun bear) and may undergo induced ovulation, although the latter does not always seem to require coitus but merely presence of a male, as is the case for the polar bear and potentially the American black bear (American black bear [1-7], Asian black bear [4], Japanese black bear [8, 9], Formosan black bear [10], polar bear [5, 11, 12], (Hokkaido) brown bear [13-16] and the sun bear [17-19]). Almost all bear species, including the giant panda, are thus seasonal breeders [14], except for certain (sub)tropical bear (sub)species of which the sun bear is probably the best studied one. In the latter bear, obligate pseudopregnancy occurs, but delayed implantation has not been described [17]. In bear species undergoing both delayed implantation and pseudopregnancy, ovulation is typically followed by a biphasic progesterone profile. This biphasic profile starts with a relatively long period of modestly elevated progesterone concentrations originating from sustained low steroidogenic activity after suppression of luteal activity during the corpus luteum (CL) dormancy phase (CLD; primary progesterone rise). A shorter second period of more marked elevated progesterone concentrations results from CL reactivation during the active luteal phase (AL; secondary progesterone rise). A similar biphasic profile has previously also been observed in giant pandas [20].

The classification of the giant panda as a bear species was not always straightforward. The herbivorous nature of the giant panda, carrying the phenotypical appearance of a real bear, historically intrigued many scientists. Carnivore phylogeny, including the position of the giant panda, is a well-documented scientific research domain that has resulted in contradictory conclusions [21-25]. First, the discussion to where to place it within the phylogenetic tree was not black and white, as it took about a century from its discovery by Armand David in 1869 until a consensus was reached and the giant panda was classified as a true bear [22, 23, 26, 27]. The giant panda is not the only true bear species to successfully transition to a vegetarian diet, but its propensity for a diet based on 99% indigestible, fiber-rich bamboo is unique. Based on the molecular study performed by Agnarsson et al. (2010), *Caniformia carnivora* split into the Canidae, containing both the family of Ailuridae (red panda) and Canidae (e.g. wolves, foxes, domestic dog), and the other caniforms [21]. Within the latter group, the Ursidae were placed sister to the Pinnipedia comprising the Odobenidae (e.g. walrus), Phocidae (e.g. seals) and Otariidae (e.g. fur seals and sea lions). In this model, the Ursidae/Pinnipedia clade was then sister to the Mustelidae. The above interrelationships are important to highlight since both the family of the Pinnipedia and Mustelidae contain species with delayed implantation and may thus be excellent examples to generate hypotheses or to extrapolate knowledge from, since studies on the exact mechanisms behind embryonal diapause and pseudopregnancy are so far largely unavailable in bear species. Nevertheless, the position of the Ursid clade against the Pinnipedia and Mustelidae sister clades in the phylogenetic tree varies critically depending on the use of a morphological or molecular classification model. The correct definition and exact time of branching of these positions, which are an indication of evolutionary relatedness, may offer a powerful tool to be used for comparative biology, consequently informing conservation strategies [21]. This is particularly true in giant pandas due to the limited access to reproductive tissues, body fluids and imaging data generated under invasive experimental conditions. Endocrinological records, ultrasound examinations and necropsies on hunted or captive bears established current knowledge on delayed implantation, CL metabolism, embryonal/fetal development and related changes in the female reproductive tract [4, 15, 28-32]. To date however, little is known about the specific mechanisms underpinning biphasic progesterone profiles representing diapause and (pseudo)pregnancy in bear species, and giant pandas in particular.

The inaccessibility to invasive biological samples from giant pandas is indeed a major limitation when researching their reproductive biology. In this study, we therefore aim to illuminate giant panda reproductive biology through available non-invasively collected samples and data. We have matched contemporary knowledge on the biphasic progesterone profile in other (bear) species with information generated after intensive reproductive monitoring of female giant pandas by reproductive endocrine profiling. Our primary goal was to define pregnancy length and thereby hypothesize on potential mechanisms behind duration of delay until CL reactivation.

## Material and methods

### Animals

Six zoological institutions provided metabolic data and urine samples for the reproductive monitoring of their resident female giant pandas for a total of 19 cycles of 6 pandas (5 pregnancies, 8 non-pregnancies and 6 pseudopregnancies) (Table 3).

Each female giant panda was housed near 1 male giant panda, in most cases allowing auditory, olfactory and visual contact. Physical contact was allowed in most cases during the time of estrus. The giant pandas were accommodated based on best practice guidelines for animal husbandry as well as recommendations given by the respective supporting Chinese giant panda experts. Giant pandas had *ad libitum* access to water and were fed a diet consisting of mainly bamboo supplemented with protein rich cake or biscuits and apples and carrots.

The pandas had free access to an indoor (ambient humidity; average temperature 18 °C) and outdoor enclosure. Specific details on husbandry and diet can be consulted in supplementary Table 7.

### Non-invasively obtained records of metabolic factors

For all 6 zoological institutions, daily fecal output was recorded by weighing the amount of feces produced, and values were disclosed at least for the duration of the reproductive cycle. Body weighing was performed during routine monitoring of the animal’s health (zoo management), such data being retrospectively acquired for our study.

Body weight was recorded on a daily basis for SB884 and SB569 and on approximately a weekly basis for SB723, SB868 and SB941. No body weight records were available for SB741. In supplementary Tables 1 and 2, information is provided on sample sizes.

**Table 1:**
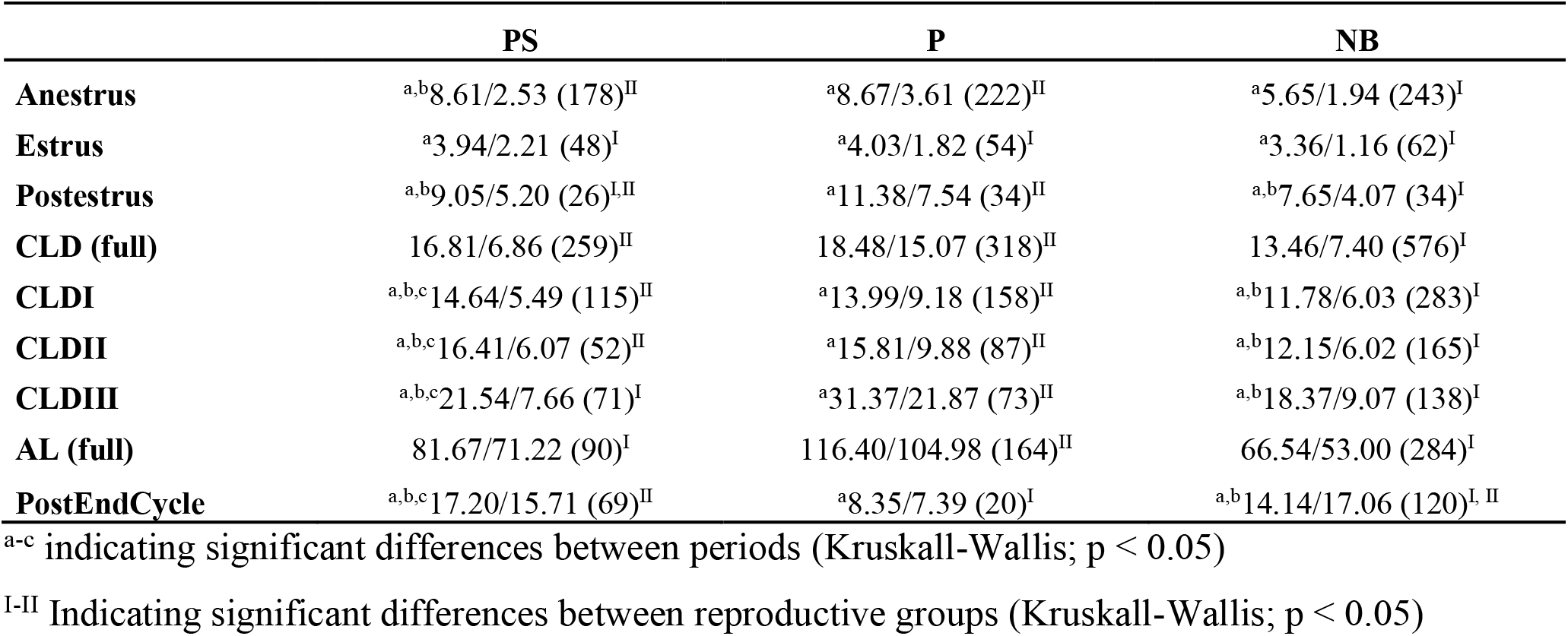
Progesterone values in ng/mL urine USpG corrected per period in cycle. Comparisons were based on Kruskall-Wallis with post hoc Dunn’s test, with significance if p < 0.05. Mean and standard deviation are listed since – within a certain period of cycling – data were normally distributed. Mean/standard deviation (number of <u>samples). CLD = CLD phase; AL = active luteal phase; PS = pseudopregnant; P = pregnant; NB = non-birth.</u>

**Table 2:** An overview of body weight and CLD details for each individual female giant panda averaged based on the included cycles. Mean (sd). BW = body weight

| Animal | Latitude | # cycles | Anestrus<br>BW (kg) | Postestrus<br>BW (kg) | ▲ BW<br>(kg) | CLDI<br>(days) | CLDII<br>(days) | CLDIII<br>(days) | CLD<br>(days) |
| --- | --- | --- | --- | --- | --- | --- | --- | --- | --- |
| SB569 | 55.95°N | 6 | 104.56 | 99.26 | 4.3 | 56.50 | 23.17 | 21.17 | 100.83 |
|  |  |  | (4.96) | (5.30) | (1.20) | (14.64) | (8.84) | (3.66) | (10.17) |
| SB723 | 47.59°N | 3* | 95.96 | 90.17 | 5.80 | 39.00 | 22.50 | 18.00 | 79.50 |
|  |  |  | (3.97) | (3.82) | (4.10) | (7.07) | (2.12) | (1.41) | (7.78) |
| SB741 | 50.60°N | 4 | 120.00 |  |  | 31.33 | 21.00 | 17.00 | 69.33 |
|  |  |  | (n.a.) | n.a. | n.a. | (8.39) | (5.29) | (2.65) | (4.62) |
| SB884 | 51.96°N | 4 | 108.14 | 101.75 | 6.39 | 18.00 | 18.67 | 20.67 | 53.25 |
|  |  |  | (1.50) | (3.22) | (2.17) | (3.00) | (3.21) | (2.89) | (9.88) |
| SB868 | 52.52°N | 1 | 81.28 | 81.00 | 0.28 | 51.00 | 26.00 | 21.00 | 98.00 |
|  |  |  | (n.a.) | (n.a.) | (n.a.) | (n.a.) | (n.a.) | (n.a.) | (n.a.) |
| SB941 | 62.55°N | 1 | 109.50 | 106.33 | 3.17 | 69.00 | 16.00 | 21.00 | 106.00 |
|  |  |  | (n.a.) | (n.a.) | (n.a.) | (n.a.) | (n.a.) | (n.a.) | (n.a.) |

\*For SB723 the atypical cycle of 2020 was excluded for determination of the CLD intervals.

### Non-invasive urine sampling

Urine was aspirated from the ground with a clean syringe and transferred to a collection vial [77]. Sample collection was pursued daily at least from pro-estrus until parturition or end of cycle.

Frequently, samples were lacking during critical periods, such as the transition from CLD phase into active luteal phase, as well as during weeks 2-3 of the active luteal phase. In pregnant cycles, sampling was often interrupted after birth due to practical limitations. During the pro-estrus and the presumed prepartum period, numerous samples were gathered throughout the day and immediately analyzed at the zoos’ facilities. In such cases, daily averages were used to generate single representative outcomes. All other samples were first stored at -20°C after sampling until analysis at the laboratories (see below).

All animal-related work has been conducted according to relevant national and international guidelines and no specific ethical approval was mandated for this study.

### Non-invasive urinary endocrine monitoring

Urinary estrogens and progesterone metabolite concentrations were determined after normalization by Urinary Specific Gravity (USpG). Using USpG-based normalization as opposed to conventional creatinine correction was advantageous since creatinine correction was previously shown to impact endocrine profiles and lead to ‘over-correction’ that was not uniform, since in turn creatinine concentrations are heavily influenced by metabolic variability. In this regard, USpG does not suffer from such metabolic variability. In a former study validating USpG-normalization as an alternative to creatinine, we clearly demonstrated significant deviation between urinary longitudinal profiles corrected by USpG *vs*. creatine corrected data during estrus and in the two to three weeks prior to onset of the active luteal phase (as defined by Wauters et al., 2018). As an artefactual result, creatinine-corrected progesterone levels showed an earlier increase because of extremely low creatinine values correlated to weight gain and reduced activity, and thus altered muscle tissue metabolism, during late corpus luteal dormancy phase. Since the active luteal phase is associated with acute, dramatic metabolic shifts, USpG-corrected estrogen profiles – but not creatinine-corrected - during late active luteal phase were discriminative for pregnancy and indicative for feto-placental involvement [78].

Urine samples of SB569 and reproductive years 2017 and 2018 of SB884 were analyzed at the MRC Centre for Reproductive Health, University of Edinburgh, Edinburgh, United Kingdom. Samples of SB741, SB723, SB884 (2019, 2020) and SB941 were analyzed at the Laboratory of Integrative Metabolomics, Ghent University, Belgium, whereas samples of SB868 were partially analyzed by the latter lab and largely by the Department of Reproduction Biology, Leibniz Institute for Zoo and Wildlife Research, Germany, rigorously adhering to the methodologies and guidelines.

For SB569 and SB884 (2017, 2018), estrogens were determined by the DetectX^®^ Estrone-3-glucuronide (E1G) Enzyme Immunoassay Kit (K036-H5, Arbor Assays™, Ann Arbor, Michigan, USA), while for the other cycles the DetectX^®^ Estrone Enzyme Immunoassay Kit (K031-H1, Arbor Assays™, Ann Arbor, Michigan, USA) was used with estrone-3-sulphate (E1S) as standard [77]. Wilson et al. (2019) described a conversion factor of 2.62, which was used in this study to convert E1G values into E1S-estimates [78].

For SB884 2017 (pseudopregnancy), an in-house progesterone test was applied compared to the DetectX® progesterone Enzyme Immunoassay Kit (K025-H1, Arbor Assays™, Ann Arbor, Michigan, USA) used for all other cycles. Therefore, progesterone levels for this cycle were excluded for further evaluation.

### Data processing

#### Definition of cycles

In this study, 19 cycles from 6 females (Table 3) were included, of which 5 cycles resulted in birth (pregnant), while for 6 cycles ovulation was not followed by mating or insemination (pseudopregnant) and the remaining 8 cycles showed no birth after ovulation and subsequent artificial insemination (non-birth, which may represent either pseudopregnancy or a lost pregnancy) (Table 3).

**Table 3:**
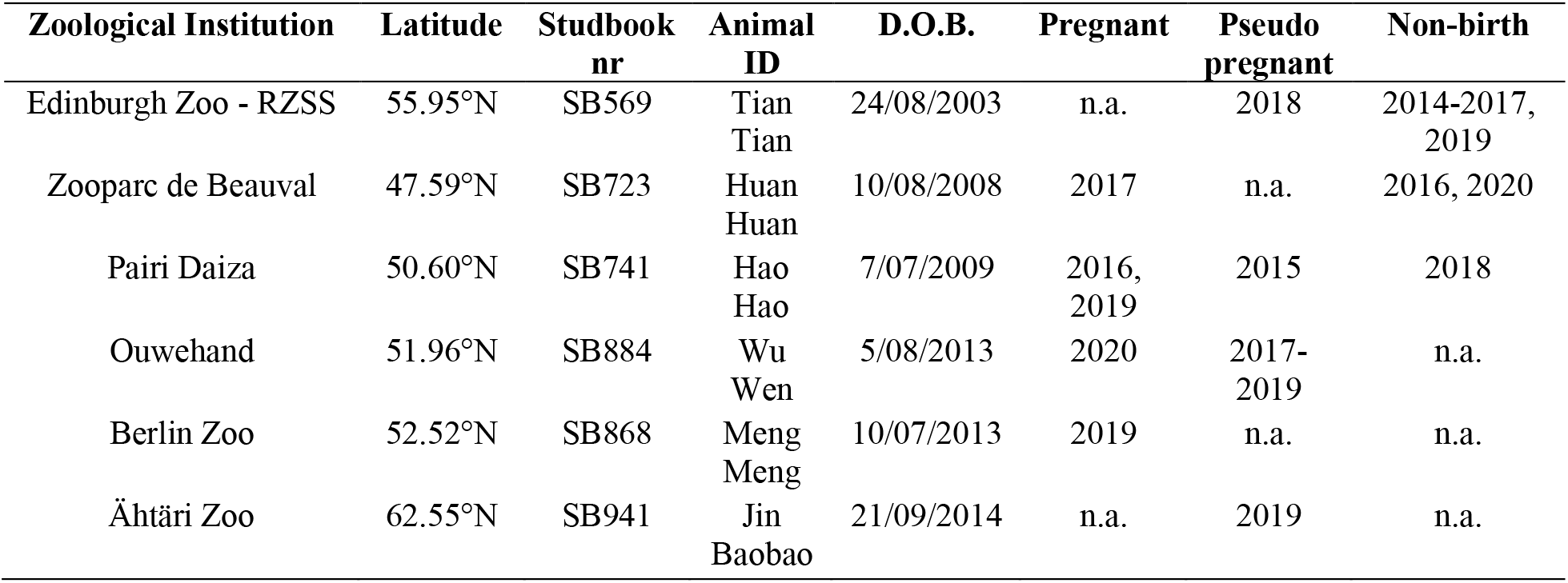
Population details on the female giant pandas included in this study (D.O.B. = date of birth). (n.a. = not applicable)

| Zoological Institution | Latitude | Studbook nr | Animal ID | D.O.B. | Pregnant | Pseudo pregnant | Non-birth |
| --- | --- | --- | --- | --- | --- | --- | --- |
| Edinburgh Zoo - RZSS | 55.95°N | SB569 | Tian<br>Tian | 24/08/2003 | n.a. | 2018 | 2014-2017,<br>2019 |
| Zooparc de Beauval | 47.59°N | SB723 | Huan<br>Huan | 10/08/2008 | 2017 | n.a. | 2016, 2020 |
| Pairi Daiza | 50.60°N | SB741 | Hao<br>Hao | 7/07/2009 | 2016,<br>2019 | 2015 | 2018 |
| Ouwehand | 51.96°N | SB884 | Wu<br>Wen | 5/08/2013 | 2020 | 2017-2019 | n.a. |
| Berlin Zoo | 52.52°N | SB868 | Meng<br>Meng | 10/07/2013 | 2019 | n.a. | n.a. |
| Ähtäri Zoo | 62.55°N | SB941 | Jin<br>Baobao | 21/09/2014 | n.a. | 2019 | n.a. |

#### Definition of subperiods based on the reproductive cycle

The period of investigation for each cycle was divided into at least 6 subperiods: anestrus (start of observations until cross over between progesterone and estrogens; [77]), estrus (cross-over until day of peak estrogen), postestrus (7 days following day of peak estrogen), CLD phase (CLD; from 7 days after day of peak estrogen until start of active luteal phase), active luteal phase (AL; from early onset of consistent increase in progesterone until end-of-cycle) and post-end-of-cycle (from end of active luteal phase until max. 30 days later). The 7-day duration of the postestrus period corresponds to the period of development from zygote to blastocyst prior to the onset of diapause, based upon knowledge in other species [79]. Since this study aimed to catalogue and comprehend the CLD phase (CLD) and its transition into the active luteal phase, the CLD was further subdivided. Based on profile observations, CLD I (plateau profile with varying length), CLD II (slow linear increase in progesterone approximately 3 months prior to birth) and CLD III (clear step-wise increase in progesterone approximately two months prior to end-of-cycle and 2-3 weeks prior to onset of active luteal phase) were defined. An additional objective of this study was to determine the transition into the active luteal phase to permit a more accurate description of pregnancy length, based on observations in the 5 pregnant cycles. For these cycles, D0 was defined by the day of parturition (Fig. 7). Based on descriptive data assessment, the early start of the transition into the active luteal phase was set at D-42/D-41, identified as the lowest point of progesterone concentration prior to onset of consistent increase. A second point of confirmation was D-37/D-36, corresponding to the start of a clearly accelerated increase in progesterone concentrations towards peak (D-20/D-19) in all pregnant cycles (Fig. 7). This second D-37/D-36 point of confirmation typically matched with the previously reported chosen formula by Kersey et al. (2010) with arbitrarily calculated values of onset of secondary progesterone rise (active luteal phase) [20].

In pseudopregnant and non-birth cycles, the definition of D0 is disputable. Additionally, frequently lacking samples sometimes made it challenging to correctly identify the D-42/D-41 and the D-37/D-36 confirmation points, particularly in pseudopregnant cycles. For those cycles, the most complete dataset was mostly the fecal output, which was used to to define timings aligned to pregnant cycle data sets.

#### Data analysis

In terms of data analysis, it is critical to acknowledge the unequal distribution of specific giant panda cycles into certain groups of interest, e.g. the pregnant (2/5 cycles from SB741), pseudopregnant (3/6 cycles from SB884) and particularly, non-birth cycles (5 of 8 cycles are from SB569). These outcomes are important, since magnitude and behavior of a specific metabolic or endocrine marker during the cycle may be influenced by giant panda maturity (adolescent versus adult), individual metabolic differences, acclimatization (newly arrived pandas versus established), climatological influences (within a cycle, but also between years) (personal observations). Additionally, interactions may occur between several of those factors. Therefore, the evaluation of data was by necessity descriptive. For any further statistical evaluation, the inevitable shortcoming of not being able to standardize for the abovementioned factors must be acknowledged when interpreting the results. Additionally, statistical evaluation was further compromised by the heteroscedastic character of both metabolic and endocrine markers, intrinsic to longitudinal profiles severely influenced by individual, seasonal and reproductive factors. Normality tests applied on single cycles and subdivided in periods did however match the assumptions, and therefore both mean (plus standard deviation) and median (plus range) are reported for those periods in the summarizing tables. General linear modelling on the complete datasets was however not applicable: evaluating the studentized and unstandardized residuals of the explored models for any of the observed markers clearly showed that the assumptions for model validity were not met. As a result of the heteroscedastic data, non-parametric approaches of biomarker comparisons were thus utilized comparing the defined reproductive periods after splitting the file based on the three groups of reproductive cycles. In parallel, the data were also assessed by comparing the latter groups after dividing the file according to the reproductive periods. To this extent, the Independent-Samples Kruskal-Wallis test followed by the post hoc Dunn’s test based on pairwise comparisons adjusted with a Bonferroni correction (significant if p < 0.05) was used.

All statistical analyses were performed with SPSS version 26.0 (IBM, Brussels, Belgium).

## Results and discussion

### Qualitative and quantitative assessment of data sets

For all cycles, nearly complete longitudinal profiles for fecal output were available, while body weight was only reported consistently (with some temporary exceptions) in SB569 (daily), SB723 (weekly), SB868 (weekly until CLD phase), SB884 (daily until active luteal phase) and SB941 (weekly). Regarding the endocrine profile, sampling efficiency largely depended on cycle category (pregnant, pseudopregnant, non-birth).

Five pregnant cycles were included in this study from 4 female giant pandas: SB741 (2016), SB723 and SB868 had daily records for all endocrine markers, while some larger sample gaps existed in SB741 (2019) and SB884 (2020) (Supplementary table 1 and 2).

Six pseudopregnant cycles were obtained from 4 females, with 1 female (SB884) representing 3 cycles. The latter data sets typically lacked numerous samples, in particular during the active luteal phase. The most complete profiles were obtained from SB884 (2018 and 2019) and SB941. SB741 (2015) and SB884 (2017) were the most poorly populated cycles, with progesterone values discarded for SB884 since a different progesterone-metabolite assay was used. For the latter two cycles, the compromised profile made it impossible to define subdivisions in the CLD phase.

Eight cycles derived from 3 female pandas were included in the non-birth category. SB569 delivered 5 relatively complete profiles with samples typically lacking during the active luteal phase and during the final 30 days of monitoring, while SB723 generated two non-birth profiles. The dominance of SB569 within this category is important for further data analysis, in combination with its atypical metabolic profiles (Wauters et al., 2022; in preparation). However, our main focus was to compare clear-cut contrasting groups i.e. pregnant versus pseudopregnant cycles.

Detailed information on sample distribution and descriptive values (mean, standard deviation, median and range) across categories of reproductive periods can be consulted in Supplementary tables 1-4.

### Defining gestation length in giant pandas

#### Alignment of pseudopregnant and non-birth cycles

The reference point for data alignment was set at D0, determined in the 5 pregnant cycles as the day of birth (Fig. 2).

**Fig 2:**
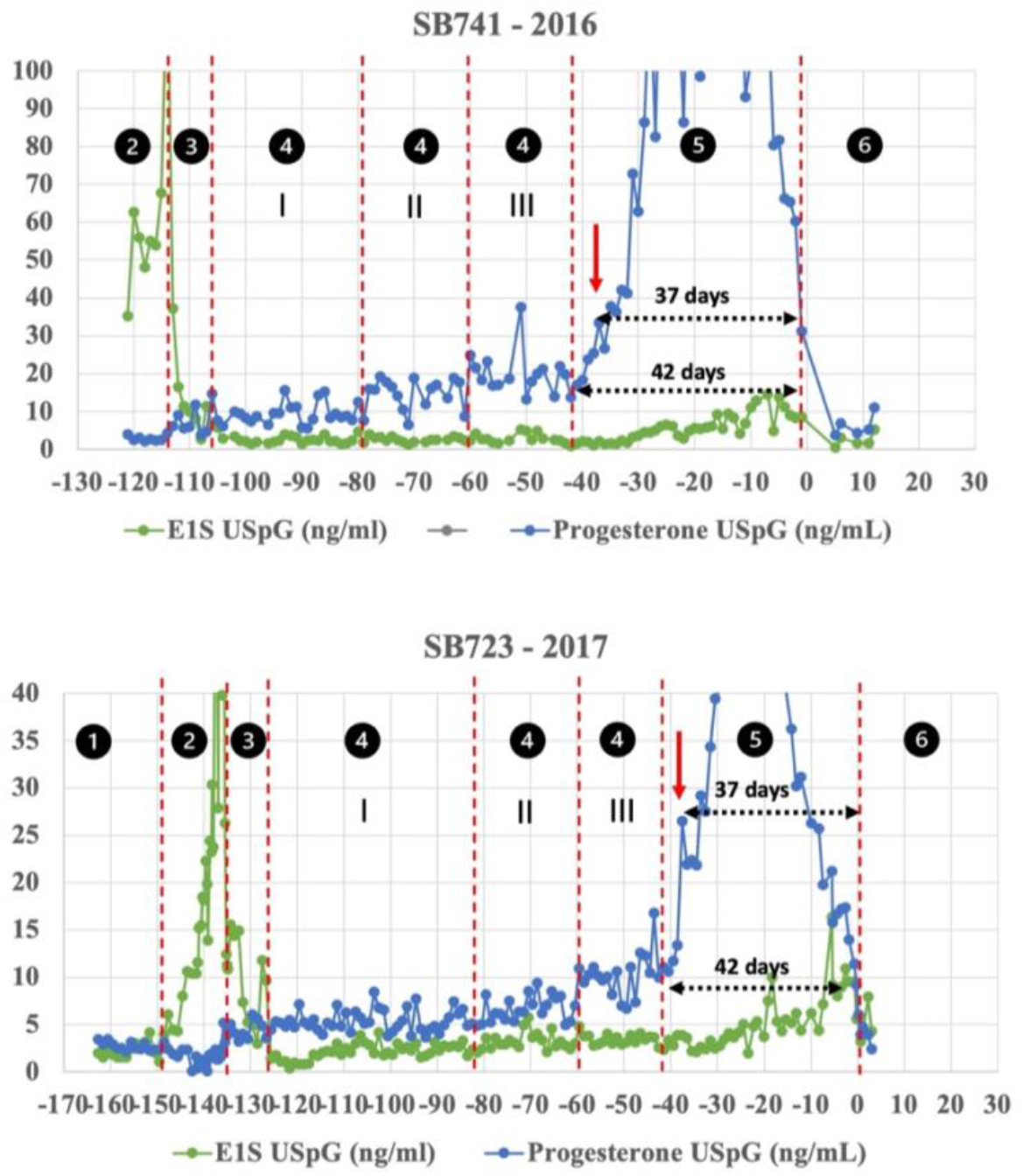
Overview of the different periods of interest in the female giant panda reproductive cycle, based on 2 pregnant cycles. 1 = anestrus; 2 = estrus; 3 = postestrus; 4 = CLD (CLD I, II and III); 5 = active luteal phase (AL); 6 = post-end-of-cycle

This definition could however not be adopted for pseudopregnant and non-birth cycles. Longitudinal monitoring of several giant pandas during consecutive cycles demonstrated that the early stage of active luteal phase was typically accompanied by distinct changes in fecal output resulting from a sudden decrease in activity, limiting time investment in bamboo intake with congruent increased progesterone concentrations. This was surprising, since progesterone was associated with appetite enhancing properties in other species [33-35]. A closer investigation of the daily fecal output numbers in pregnant cycles leads us to the definition of an alignment procedure for all non-pregnant cycles (Fig. 3). More specifically, 2 reproducible points of recognition were identified. In pregnant cycles, daily fecal output was typically at half-maximum 30.10 ± 1.43 days prior to birth (= one week after the onset of accelerated increase in progesterone), remaining around the same level for 1 to 3 days, prior to a further lower plateauing (1 to 3 days) point at 27.60 ± 1.50 days prior to birth (Fig. 3A). In pregnant cycles, fecal output numbers then typically continued to decline, whereas stabilization of fecal output numbers was observed in non-pregnant cycles (Fig. 3B&C). For this reason, in non-pregnant cycles, D-28 was matched with the ‘lowest’ point of fecal output. Based on this alignment, half-maximum numbers were observed at 30.50 ± 0.50 and 30.44 ± 0.63 in non-birth and pseudopregnant profiles, respectively, which corresponded to the pregnant cycles (Fig. 4). In only one non-birth cycle (SB723 2020), the D-28 criterium was hard to match, most likely because of the seasonal impact on this atypical cycle (estrus occurred in December and cycle length was significantly shorter compared to previous cycles and ending in March 2020).

**Fig 3:**
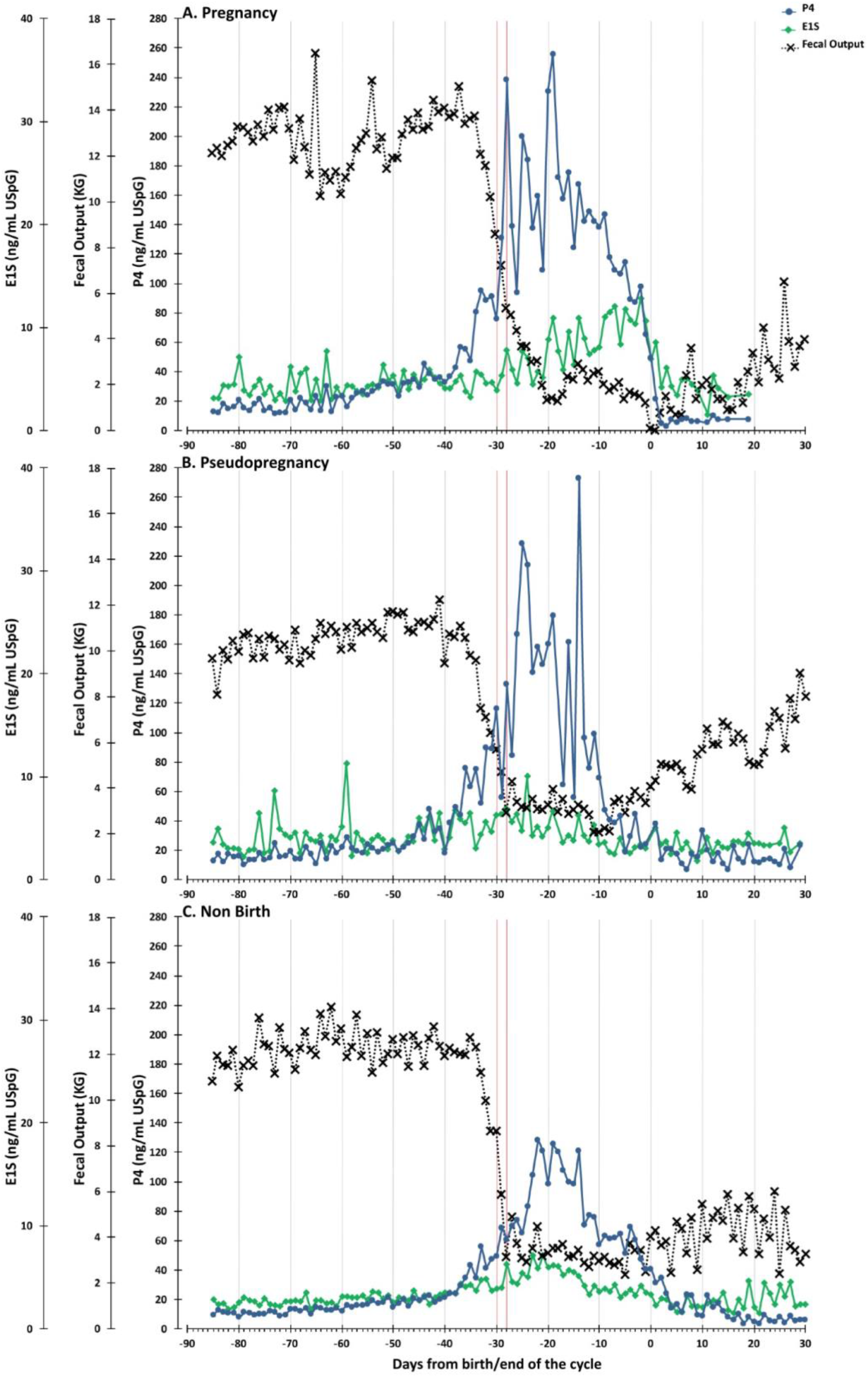
Average urinary estrogen and progesterone metabolite profiles, average faecal output profiles for pregnant, pseudopregnant and non-birth cycles in female giant pandas.

**Fig 4:**
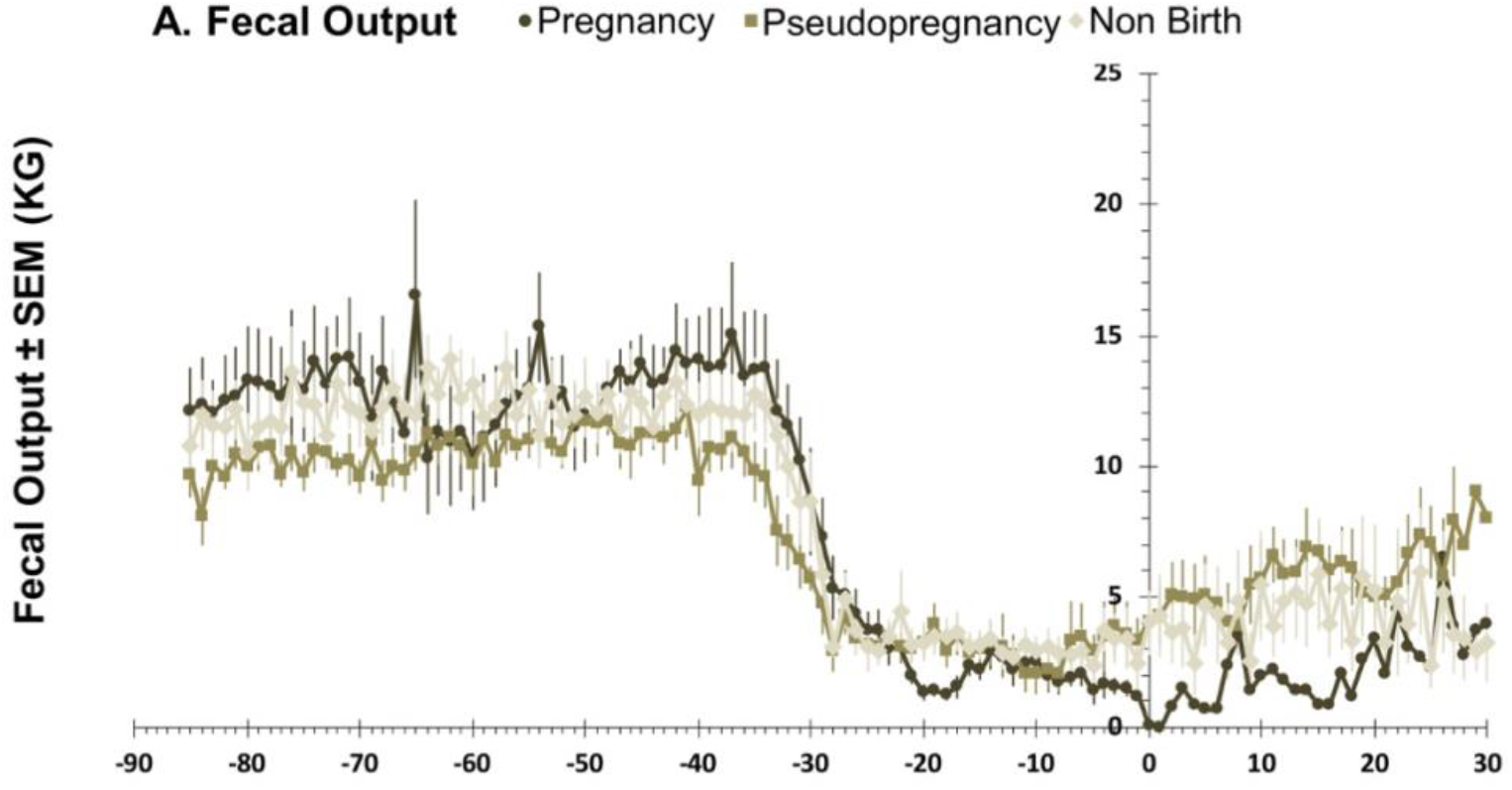
Fecal output profiles for pregnant female giant pandas. Utilising alignment of the faecal output profiles in pregnant cycles, two points of recognition were chosen to align pseudopregnant and non-birth cycles with respect to D0 (= end-of-cycle).

#### Reactivation of CLs 42 days prior to birth: pregnancy takes 37-42 days in giant pandas

As described above, we used a quantitative metabolic marker (fecal output) to align all cycles. Other bear studies have similarly used alternative non-invasively obtained, non-endocrine markers (for example temperature and activity data) to estimate implantation and birth date during hibernation [36, 37].

Herein, decreased fecal output due to reduced intake of nutrients as a consequence of limiting activities and thus lowered basal metabolism in favor of reproduction, occurred within one week following the progesterone increase, known as the start of the secondary progesterone increase. We confirmed distinct changes, exclusively monitored in the USpG-corrected progesterone profile, from D-42 onwards, with D-42 representing a nadir in progesterone prior to consistent increase, the latter being accelerated and becoming statistically significant from the averaged primary rise values, typically at D-37 (Fig. 2). Based on this information, we suggest gestation in giant pandas to be of fixed duration, taking not more than 42/43 (D-42) days from early reactivation of the CL leading to significant luteal progesterone production supporting embryonal growth from 37-38 (D-37) days before the birth of an altricial cub.

A pregnancy length of maximum 42 days is significantly shorter than reported for any of the other 7 bear species. Exact information on other bear species’ CL dynamics and pregnancy durations is extremely limited and restricted to estimations of potential timing of CL reactivation (approximately 2-4 weeks prior to implantation) and embryonal implantation (60 days prior to birth) [2, 5, 16, 31, 32, 36]. In the brown bear, the mean date of implantation was documented to be the 1^st^ of December (standard deviation (sd) = 12 days), with subsequent parturition around 26^th^ of January (sd = 12 days) and thus average duration of post-implantation gestation of 56 days (sd = 2 days) [36]. Based on the scarce information available in other bear species, pregnancy – including pre-implantation events such as development of the embryonic folds – lasts approximately 70-88 days and is thus at least 4 weeks longer compared to the gestation length defined in this study for giant pandas. The shortening of pregnancy by almost one month may explain the findings of Griffiths et al. (2015), who reported that transition from colostrum to main phase lactation only took place approximately 30 days post-partum in giant pandas [38]. The validity of this short pregnancy model, particularly the exact pregnancy duration of 42 days, is additionally supported by the study of Li and Smith (2020) [39]. These authors investigated ossification and skeletal anatomy in neonatal ursid cubs and concluded that other bear species are not exceptionally altricial relative to other caniforms, despite the impressive difference in body size between newborns and adults. In contrast, for the two newborn giant panda cubs investigated in this study, skeletal maturity levels matched up to beagle fetuses of only 42-45 days old, corresponding to 70% of the total gestation period. In addition, in canine species, skin pigmentation may start to develop at 37 days onwards, but is only noticeable over the whole body from day 46 onwards. This is in accordance with the development of the black and white patches in giant pandas during the first week post-partum. Body hair presence is evident between 40 and 46 days of development in dogs, aligning with the physical appearances of giant panda cubs at birth [40]. A pregnancy of 37-42 days is thus scientifically plausible for giant pandas.

For the finetuning of the definition of pregnancy length in other bear species, longitudinal non-invasive endocrinology studies should be launched. Urinary profiles corrected with USpG rather than creatinine may support a similar unraveling of the reproductive biology in other bear species, and as such aid to determine exact pregnancy length as well as proper investigation into the biphasic progesterone rise. Nevertheless, obtaining non-invasive matrices such as urine or feces during hibernation is fairly unfeasible and therefore explaining the limited resources of information in these species up until today.

### The CLD phase redefined

#### Tri-phasic increase in progesterone

In giant pandas, the CLD phase is often described by a modest and long-term primary increase in progesterone, commencing shortly before peak estrogen secretion during estrus [41, 42]. This primary increase is described to last for a period of 2 to 4 months [20], until the start of a secondary more obvious increase in progesterone following CL (re)activation (active luteal phase) (Fig. 2 & 3). This biphasic character is also reported for many of the other bear species, indistinguishable between pregnant and pseudopregnant cycles [2, 5, 16, 43]. The mechanisms behind this biphasic increase remain obscure but are associated with a previous suppression of CL activity allowing an embryonal diapause, maintained by low progesterone levels, in conceived animals. It is generally accepted that delayed implantation or embryonic diapause serves to disconnect the period of mating from the period of birth, facilitating lactation and/or raising cubs under the most favorable environmental conditions [44]. The pathways involved may perhaps be partly extrapolated from the best studied carnivorous delayed implanter, the mink. Delayed implantation in mink additionally serves to synchronize embryo implantation/development resulting from several follicular phases and matings [45], an elegant strategy to increase genetic diversity and population fitness if matings originate from different paternity. This phenomenon has also been described for the black bear and presumably also exists in other bear species such as polar bears [3, 46]. In the mink, CLs of a previously induced ovulation undergo involution but continue to secrete steroids at concentrations corresponding to the primary rise in progesterone, insufficient to support implantation or development of embryos (+- 8 ng/mL in plasma) [45]. Follicle development continues with potential ovulation every 6-7 days allowing mixed paternity during one reproductive cycle. The presence of multiple underdeveloped follicles on ovaries of female giant pandas observed post-ovulation during AI procedures may therefore be a remnant of their ancestor’s strategy to maintain population fitness, abandoned during further evolutionary development from carnivore to herbivore when this was no longer matching their metabolic capacities. Indeed, the cost of reproduction, e.g. by synthesis of steroids and their subsequent metabolic actions, is high and a polyestrous strategy may thus have been abandoned [47, 48].

Steroidogenic capacity of the CL during the delay phase was indeed also confirmed in the Hokkaido black bear [49] and is presumably also responsible for the primary progesterone rise described in other bear species, including the giant panda. The biphasic profile of progesterone secretion thus results from involuted CLs initially retaining their basal steroidogenic competence during the CLD phase and then further developing during the active luteal phase by advanced luteinization [50]. From the current dataset, we could however derive that the CLD phase in giant pandas is not a static period of long-term slightly elevated progesterone secretion at stable levels. More specifically, we observed three periods with a first turn-over point characterized by ‘bending’ of the progesterone curve towards higher levels (Fig. 2; Fig. 5). Though the definition of this event is only based on rather subjective observations, potentially explaining the relatively wide range of onset, this was observed in all pregnant cycles at D-81.20 +- 5.85 (range D-89 to D-73) prior to birth (= CLD II), i.e. a relatively fixed onset in relation to the date of birth, but with a more variable length from the point of ovulation for each giant panda and cycle (e.g. 23 to 59 days post peak estrogen) (Supplementary table 5). A more distinct and significant step-wise increase in progesterone is further monitored at 60.80 +- 3.83 days (range D-63 to D-56) prior to birth (= CLD III), again characterized by a relative fixed time point if related to birth. These CLD turnover points are also present in the pseudopregnant and non-birth cycles, at reasonably similar timings. In these groups, progesterone starts shifting slowly towards slightly higher levels at 82.25 +- 5.68 (range D-88 to D-76) and 83.75 +- 9.35 (range D-96 to D-70) respectively, while the more abrupt step-wise increase was recorded 64.75 +- 3.50 days (range D-69 to D-61) prior to the presumed end-of-cycle in pseudopregnant cycles and 61.13 +- 2.59 days (range D-65 to D-57) in non-birth cycles. On average, dormancy II and III each lasted 2-3 weeks, with the most consistent interval length for CLD III (resp. 21.06 +- 6.30 and 19.59 +- 3.24 days) (Supplementary Table 5). This step-wise increase in progesterone levels occurring approximately 2 months prior to the theoretical end-of-cycle was more pronounced compared to the previous stages of the CLD phase in all cycles. Moreover, this increase showed to be more distinct in pregnant cycles with significantly higher progesterone concentrations compared to pseudopregnant and non-birth cycles (Table 1). Interestingly, the latter observations were also true for estrogens during this last stage of the CLD phase, indicating that both steroids were secreted by the same endocrine organ and resulting from the same trigger. In some species, CLs are known to produce estrogens. Other related delayed implanters such as species from the Phocidae (e.g. spotted seal) and Ursidae (e.g. Japanese black bear) display some capacity of estrogen synthesis by the CL, predominantly in the early stages of reactivation [51, 52]. The increase in steroids at this phase of cycle in giant pandas is thus likely resulting from an increasing CL activity. While the timing of onset is not different between pregnant and pseudopregnant cycles, the significant higher levels of these sex steroids from around D-60 onwards in pregnant cycles may perhaps indicate active signaling between blastocysts and mother tissues after an initial luteal increase in progesterone, enhancing overall progesterone production and thus facilitating the survival and reactivation of the blastocyst due to increased availability of histotrophic substances and other regulating factors. Remarkably, this significant increase in steroids that are presumably luteal two months prior to birth/end-of-cycle aligns with the pending implantation described in all other bear species for which a post-implantation development of approximately 60 days is reported [4, 9, 31]. In addition, the upward trends in P4 concentrations from on average 2 to 3 weeks before, e.g. during CLD II, are in accordance with the time of initial CL reactivation in other bear species during hibernation (Fig. 5). So could it be possible that the female giant panda has evolved a mechanism to escape from this evolutionary conserved timeline and has developed a protective mechanism to prolong the CLD phase in favor of a shorter pregnancy length/active luteal phase, driven by their metabolic limitations after the transition to low energetic bamboo as their main food source? The data generated in this study are indeed highly indicative.

**Fig. 5:**
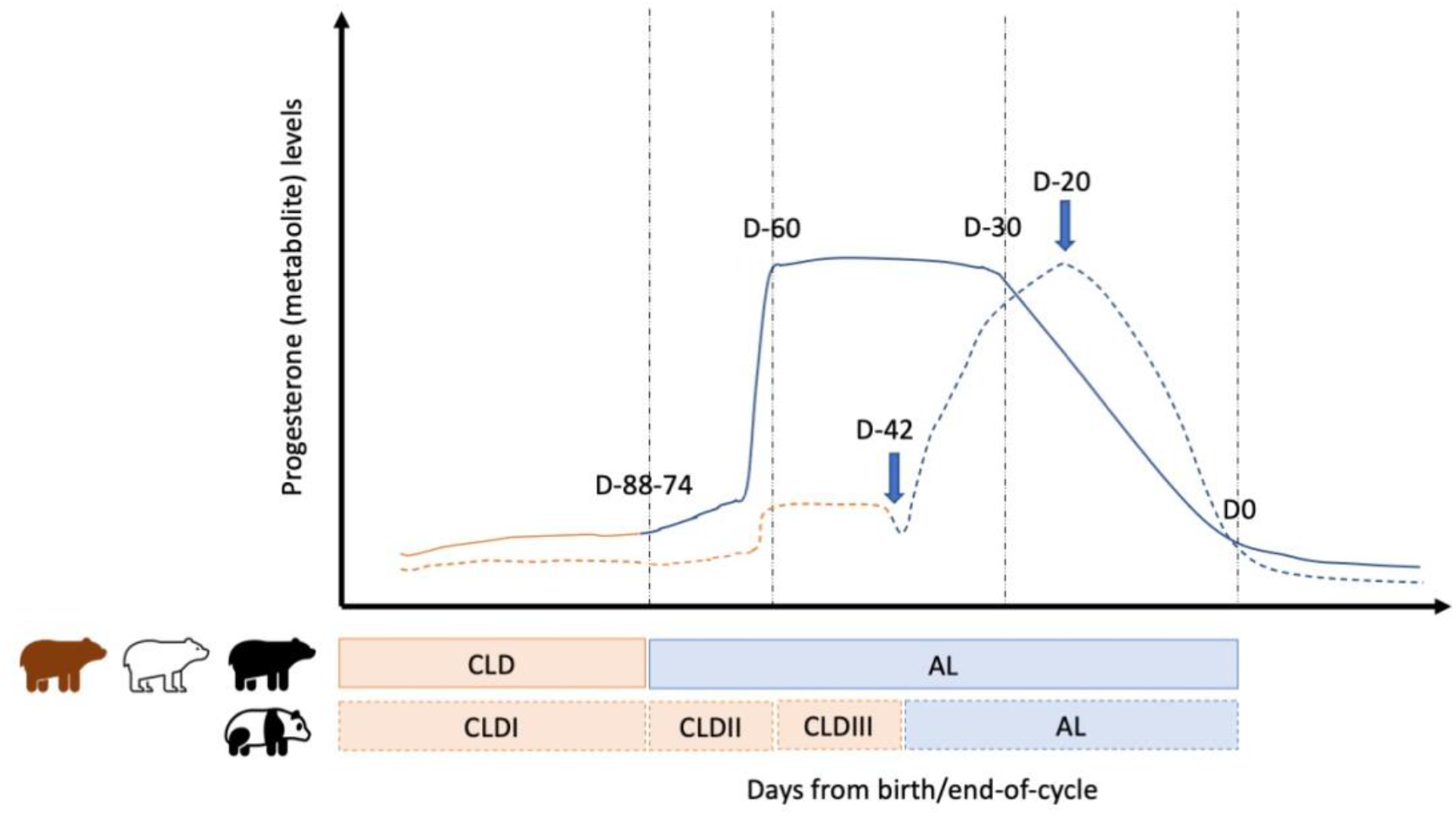
Schematic representation of the progesterone profile in bear species based on interpretations from literature (other bear species) and own data (giant panda). Giant pandas seem to extend the CLD phase beyond the evolutionary conserved timeline by suppressing CL reactivation. Together with other mechanisms at play (Wauters et al., 2022; in preparation), this ultimately results in a shorter pregnancy duration (AL) compared to the other bear species, and the birth of more altricial cubs.

#### Metabolic status of the female giant panda defines the length of stage I CLD phase: hypothetical proof

While the occurrence of the vernal equinox - by the melatonin-prolactin (PRL) cascade - is clearly the main trigger for CL reactivation and implantation in the mink, a more complex regulation seems to be in play for the escape from diapause in giant pandas and perhaps bear species in general. The latter hypothesis is strengthened by the observation that, although most pandas/cycles in this study timed estrus closely to the vernal equinox while onset of their active luteal phase was somewhat aligning with the summer solstice, it seems that higher latitude pandas (SB569 and SB941) as well as lower weight pandas (SB723 and SB868) show longer periods of delay in the CLD phase compared to lower latitude/higher weight pandas (SB741 and SB884) (Table 2, Supplementary table 6, SB569: 100.83 (mean) ±17 (sd) days (n=6), SB941 (n=1): 106 days, SB723 (n=2): 79.50 ± 7.78 days, SB868 (n=1): 98 days, SB741 (n=3): 69.33 ± 4.62 days, SB884 (n=3): 53.25 ± 9.88 days). In other words, although Thom et al. (2004) reported no significant correlation between length of delay and latitude (studied on the species level and thus not on the individual level within a species) [53], it seems that colder temperatures and smaller-sized pandas may need more time to prepare themselves for the energy demanding processes of CL reactivation, implantation, embryonal growth and lactation. The conversion from carnivore to herbivore may have skewed the ancient well-conserved largely photoperiodic regulated form of delayed implantation still in play in other bear species. In the absence of a period of hyperphagia and/or hibernation metabolically supporting a successful pregnancy [14, 54-56], giant pandas seem to have shifted to a mainly metabolic controlled timeline of embryonic diapause featuring a permissive metabolically driven decision-maker for the initiation of advanced luteal steroidogenesis and subsequent implantation. The potential physiological mechanisms behind metabolic control will be discussed below.

The understanding of central and peripheral control of reproduction is incomplete due to the complexity of the pathways involved which are intertwined with many other physiological processes such as stress and metabolic homeostasis [57]. New concepts are arising, but knowledge gaps make it challenging to complete the puzzle, particularly for species with little opportunities to study specific tissues and organs. Nevertheless, at least two highly conserved interacting pathways are described controlling reproduction efforts, which may help explain the reproductive biology in giant pandas and how they may have succeeded in customizing the onset/length of gestation.

Proper functioning of female reproductive biology involves complex interactions between the brains, pituitary and gonads. The earliest recognized pathway is the commonly known cascade in which GnRH (hereinafter referred to as GnRH I) from the hypothalamus induces the release of gonadotropins (LH, FSH and PRL) from the pars distalis of the pituitary into the circulation, inducing down-stream effects e.g. at the level of the reproductive tract. In seasonal animals this pathway is only activated during the breeding season as a result of changes in daylight, translated into differential release of melatonin [58-60]. More recently, it was recognized that the pars tuberalis of the anterior pituitary played a key role in transducing decreased melatonine-signals into a ‘command to fire’ GnRH neurons [59]. In short – for long day breeders – time-dependent binding of melatonin on its receptor of pars tuberalis’ thyrothrohps induces expression and secretion of the thyroid-stimulating hormone (TSH), which then readily reaches the adjacent median eminence of the mediobasal hypothalamus (MBH). Here, TSH will bind on its receptor expressed in ependymal cells. The latter causes increased expression of the DIO2 gene encoding the thyroid-hormone activating enzyme metabolizing thyroxine (T4) into biological active T3. In parallel, expression of DIO3 (rT3 inactive product) is suppressed. DIO2-produced T3 influences the microanatomy at the median eminence of the MBH by regulating the neuroglial interactions between GnRH nerve terminals and glial endfeet in such a way that the GnRH terminals are in close contact with the portal capillaries, facilitating seasonal GnRH secretion [58-60]. The dependence of the GnRH pathway on peripheral thyroid hormone availability in the brains may offer an explanation for the reduced libido often seen in giant pandas and may have also facilitated other pathways to play a controlling role in reproduction. Due to their dietary shift to bamboo, giant pandas developed the ability to detoxify cyanide to thiocyanate, assisted by the rhodanese-enzyme [61]. Thiocyanate is however well recognized as a thyrostatic agent [62] and thus likely responsible for reduced thyroid hormone production in giant pandas, hence potentially compromising thyroid hormone synthesis in giant pandas and consequently suppressing the commonly known gonadal GnRH pathway. In short, this may indicate that cyanide levels in the diet of giant pandas, particularly in captivity where pandas can only eat what is offered, may directly influence the reproductive biology of an individual. Investigating the potential impact of plant-derived thiocyanates on giant panda reproduction as a consequence of diet choice in captive giant pandas may thus reveal new insights in the general management of giant pandas in captivity but may also be key in the further unraveling of their reproductive biology. However, a clear indication of other pathways at play is that, even at the same latitude, and under similar environmental conditions, a reasonable spread is seen in ovulation dates for female giant pandas. The same observation is true for onset of the active luteal phase, or duration of the CLD phase, and thus potential birth season. The latter contrasts with other delayed implanters which show a more ‘synchronized’ breeding season e.g. the carnivorous mink (birth in May), other bear species (depending on the species 1-1.5 month covering period during winter/early spring) or the herbivorous caribou (births in May) [63]. Particularly in giant pandas, other factors seem to allow individual fine tuning of the reproductive cycle and onset of specific events. An interesting hypothesis to explain this, is their phylogenetic background shifting from carnivorous to herbivorous. In herbivores, parturition is often synchronized based on plant phenology [64], closely linked to seasonality and photoperiodism and thus correlated to environmental factors inducing food abundance. Nevertheless, particularly in these species, climate change has illustrated the role of individual phenotypic plasticity on reproductive traits by reducing or prolonging the pregnancy based on the female physical condition, not only correlated to the metabolic state during late gestation, but remarkable as well to the condition of the female at the start of the breeding season [63, 65]. Similar observations were reported in bear species highlighting the importance of pre-breeding or pre-denning nutritional status [54-56, 66]. The latter clearly illustrates that the duration of pregnancy can be prolonged based on metabolic triggers. Additionally, while CL reactivation in the mink is solely PRL-driven, successful implantation still requires a specific protein message, likely involving the evolutionary highly conserved glucose-6-phosphatase isomerase, which is assigned an essential role in both glycolysis and gluconeogenesis [67]. The latter is another confirmation of the high link between seasonality and metabolic state as key players in the central control of reproduction [68, 69].

One of the mechanisms involved may be the relatively unknown GnRH II pathway. Photoperiodic activation of the reproductive system is governed by the first discovered isoform of GnRH in the mammalian brain, GnRH I, regulating the release of gonadotrophins, as previously discussed, but potentially compromised in giant panda through the involvement of thyroid hormones. A more ancient and highly conserved GnRH II was more recently discovered, as were several other structural variants. GnRH II is produced in both brain and periphery and is therefore likely to cover a wide spectrum of functionalities, far beyond reproduction only. In the periphery, both GnRH I and GnRH II regulate cell proliferation and hormonal secretion from the ovary and placenta in an autocrine/paracrine way. However in the brain, they modulate reproductive behavior in different but complementary ways as GnRH II is not a strong stimulator of gonadotropin secretion and merely acts as permissive neuroendocrine regulator of female reproduction, driven by energy status [70-73]. GnRH II neurons are a subpopulation of GnRH neurons that are localized in other parts of the brain compared to GnRH I and express glucokinase mRNA, a marker for glucose sensitivity. Consequently, they decrease their firing in response to low glucose concentrations and metabolic inhibition [74]. They additionally express KATP channels that are responsive to leptin, insulin and free fatty acids and thus play a global role in energy homeostasis. It is very likely that this GnRH II pathway is involved in onset of estrus in giant pandas. It may also explain why in some pandas estrus occurs (driven by the GnRH I pathway; seasonal photoperiodic influences) without full display of behaviors (inhibited GnRH pathway caused by suboptimal metabolic state). Interestingly, Tang et al. (2019) recently demonstrated GnRHR polymorphism in the giant panda population to be correlated with differences in fertility [75]. It is therefore tempting to speculate that interacting mechanisms between the GnRH I and GnRH II pathways may be responsible for termination of phase I of the CLD phase and may play a role in delaying onset of the active luteal phase by inhibiting effects on the evolutionary conserved timing of GnRH I firing (other potential key players: kisspeptin, GnIH, leptin and other appetite regulating peptides) [57, 76]. Indeed, observations from our study indicate an important individual factor involved in onset of endocrine and metabolic changes during the CLD phase (Table 2).

### Giant pandas push back the active luteal phase: rationale behind stage II and III of the CLD phase

Most other seasonally breeding bear species go into hibernation prior to continuing their pregnancy, which includes reactivation of the CL, implantation and embryonal/fetal development (Fig. 6). Within a maximum of a few weeks into hibernation, the CLs reactivate, leading to implantation approximately 60 days prior to birth. These processes are accompanied by increasing progesterone values, first modestly, but then rather steeply around the time of implantation (Fig. 5). The onset of these events (CL reactivation and implantation) are relatively fixed within and across bear species and strongly correlated to photoperiodicity cues leading to a narrow time-framed birth season (Fig. 6) [2, 4, 5, 9-11, 13, 16, 17, 19].

**Fig 6:**
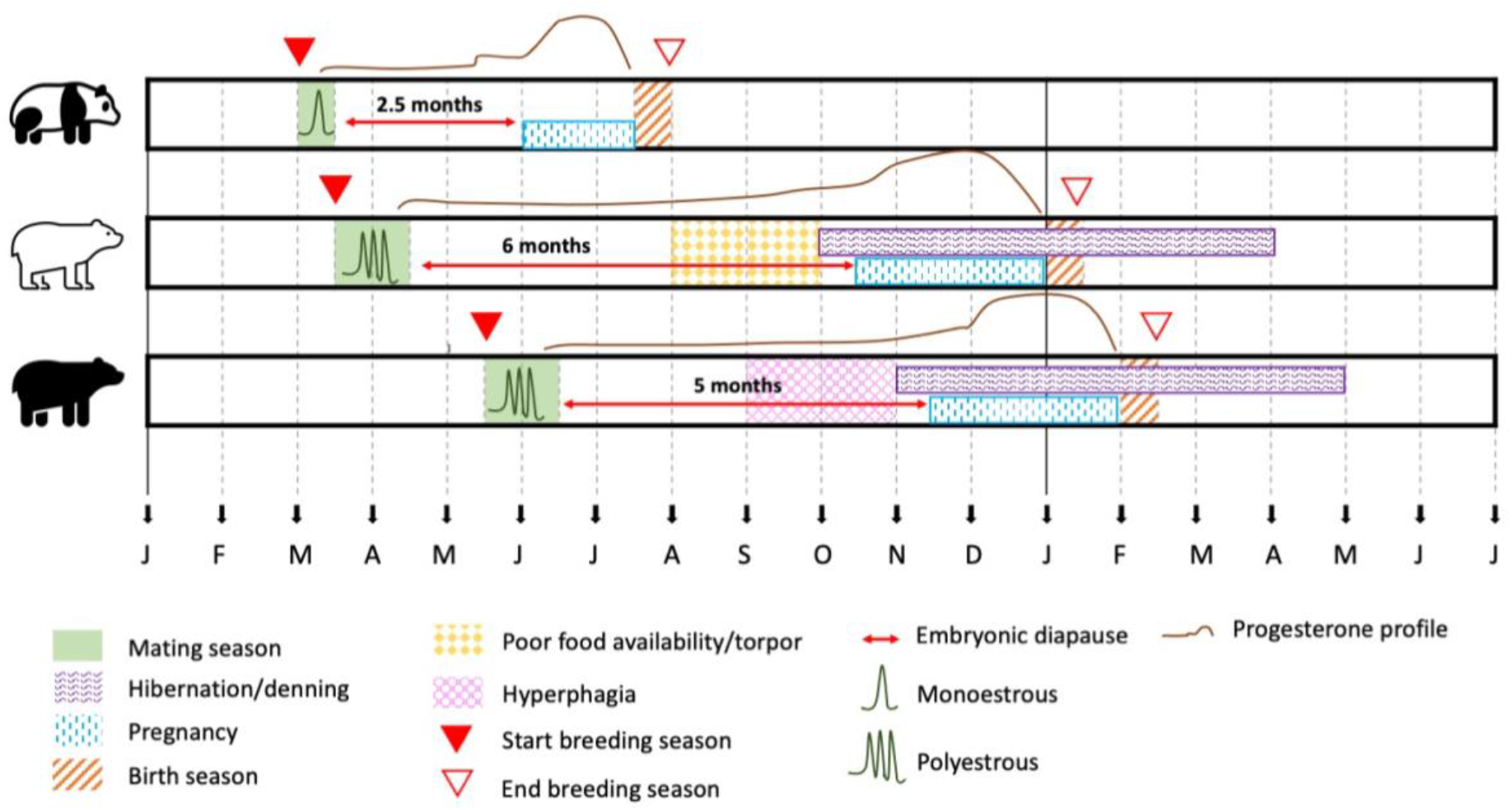
Comparative overview of the female reproductive cycle in a hypothetical average female individual for three bear species. In giant pandas the breeding season starts early and is short, with only one oestrus occurring followed by a relatively short CLD phase, which allows birth during the same year’s most favorable environmental conditions, therefore avoiding the need to hibernate. This delay until potential implantation is still mandatory to allow restoring energy reserves after the energy demanding estrus season and to prepare for potential pregnancy. After accumulation of sufficient energy reserves, pregnancy can continue, but is restricted in duration compared to the other bear species, because of the limited capacity of herbivorous giant pandas to build up energy reserves. While embryonic development of giant pandas is hypothesized to take approximately 3 weeks after implantation (total pregnancy duration 42 days), a post-implantation of approximately 60 (56) days is accepted in the other bear species (total pregnancy duration 60 days +- 2-4 weeks). Giant panda cubs are therefore much more altricial compared to other bear species.

**Fig 7:**
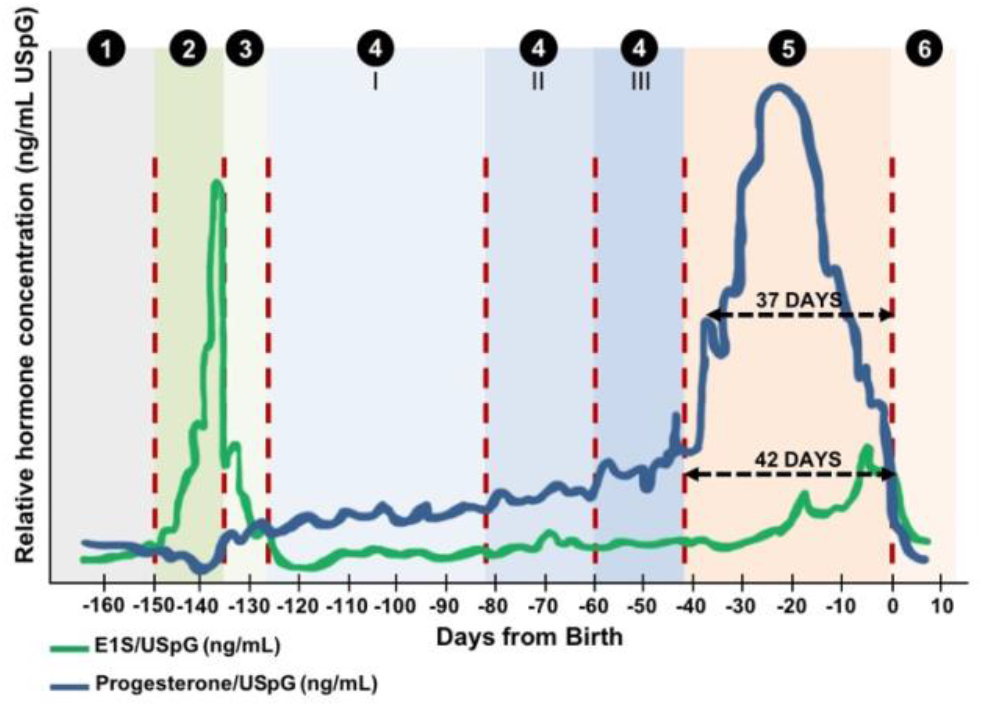
Schematic overview of the different periods of interest in the female giant panda reproductive cycle. 1 = anestrus; 2 = estrus; 3 = postestrus; 4 = CLD (CLD I, II and III); 5 = active luteal phase (AL); 6 = post-end-of-cycle

Our dataset, showing three phases in the giant panda progesterone profile, indicates that the first gradual increase in progesterone, noticed at +- D-81, may correspond to the observed upward shift in progesterone levels in other bear species at the early phase of their hibernation period (CL reactivation) (Fig. 6). Around D-60 - the timing of presumed implantation and maximum CL capacity in other species - a clear stepwise increase, significant in both pregnant and pseudopregnant females, is observed in progesterone for giant pandas (Table 1; Fig. 6).

The increasing ovarian steroids from the partly reactivated CLs around D-60 most likely support the activation of the secretory uterus to create a favorable environment for the pending rapid development of the blastocyst and its embryonic membranes. Early changes in the uterus, prior to embryonic folding, elongation and implantation, are indeed also observed in other species. This includes morphological alterations in the endometrium (folds and glands) as well as secretory activity (uterine milk) [31, 45].

Nevertheless, the further increase of progesterone to pregnancy supporting levels allowing further blastocyst and embryonic fold development and subsequent implantation, appear suppressed and delayed in giant pandas until D-42, or even D-37 and onwards. The onset of the CL reactivation in other bear species during challenging times (low temperatures, short days, reduced availability of food resources) is possible thanks to their strategy of hibernation, allowing a longer pregnancy duration and more developed bear cubs. Hibernation is obligate in most seasonal bear species and typically occurs after a period of hyperphagia allowing body weight accumulation, but facultative in the female polar bear. Non-pregnant and male polar bears usually remain active during autumn/winter but may undergo periods of torpor to reduce energy expenditure in periods with food scarcity, typically occurring from late summer onwards [11, 66]. The strategy of hibernation in female polar bears is unfortunately not always effective to safeguard pregnancy and highly dependent on their pre-denning weight and body condition since they lose around 43% of body mass during hibernation. As a result, 33% of the pregnancies are estimated to result in fetal loss [11, 54, 55, 66]. Due to their low-energy diet and dysfunctional digestive system preventing sufficient nutrient accumulation for long-term capacity, giant pandas most likely had to abandon the strategy of hibernation completely. As an alternative, giant pandas’ diapause is on the one hand shortened compared to other bear species, facilitating pregnancy and early lactation during most favorable conditions within the same half year (Spring-Summer), at times where food resources are still guaranteed. On the other hand, diapause is extended by delaying the onset of CL reactivation e.g. by shortening the energy demanding active luteal phase with high-level progesterone production.

If their reproductive cycle depended only upon availability of shoots and nutrient rich young bamboo, it would be most favorable for most pandas to continue pregnancy without any delay. Nevertheless, both the energy loss during estrus and the need to ‘refuel’ for potential pregnancy probably resulted in the obligate character of a customized delay in CL reactivation (Table 2; Supplementary table 6). During dormancy phase II and III, respectively corresponding to CL reactivation and implantation in other bear species during hibernation, reduction of energy loss in favor of fueling reproductive processes is additionally controlled by decreased activity. The activity rate is progressively decreasing during the active luteal phase, causing bamboo intake to decline significantly in the early stage of the active luteal phase, underscoring the utility of non-invasively obtained, non-endocrine markers such as fecal output number as an easy applicable indicator for having entered the final crucial phase in the giant panda reproductive cycle, i.e. potential pregnancy.

## Conclusions

The evolutionary triggers for CL reactivation appear to be conserved in giant pandas, as demonstrated by the shifts in progesterone concentrations, but giant pandas seem to have developed a strategy to customize CLD length and postpone full reactivation of the CL in favor of their compromised metabolism, as an alternative to hibernation. This results in a shorter pregnancy duration and more pronounced altricial cubs compared to the other bear species. The increases seen in estrogen and progesterone during the progression of the CLD phase, and more particular from 2 months prior to the end of cycle and approximately 2-3 weeks before the abrupt secondary increase in progesterone, likely result from pathways involved in evolutionary conserved photoperiod-steered seasonality and metabolic state monitoring, originally targeting a pregnancy of plus 2 months, which has been shortened to 37-42 days (5.5-6 weeks) in giant pandas. Although studies in other bear species indicate no increase in CL sizes during the CLD phase, novel ultrasound techniques could be applied to support current hypothesis by documenting CL sizes during the different identified stages of the CLD phase and the active luteal phase.

Based on this information, and thanks to the successful and easy alignment of all non-pregnant cycles (except one), prediction of potential birth (= end-of-cycle) can be performed accurately as soon as 6-7 weeks prior to potential birth, and less accurately around 60 days prior to birth. The latter allows zoos to prepare timely for potential birth and may prevent excessive night guarding prior to and after the presumed end-of-cycle, in both pregnant and non-pregnant cycles.

## Acknowledgements

The authors would like to express their greatest appreciation for the support offered by the giant panda experts from both the China Conservation and Research Centre for Giant Panda (CCRCGP) and the Chengdu Research Base of Giant Panda Breeding (CRBGPB), and would like to express a special gratitude towards Mrs. Cindy Luo and Mrs. Wei Ling and their respective teams for their excellent communication, hospitality and administrative services.

The authors would also like to thank all participating zoological institutes with special gratitude towards the keepers involved in sample collection.

The authors would also like to thank Dirk Stockx, Tom Cools, Mareen Albrecht and Katrin Paschmionka for the assay assistance at Ghent University and the Leibniz Institute for Zoo and Wildlife Research. This research is financially supported by the Pairi Daiza Foundation, Ghent University and Beauval Nature.

