## Supplementary files for "Evolutionary survival strategies of the female giant panda: optimizing energy resources and expenditure prior to pregnancy by postponing corpus luteum reactivation"

### 1 Supplementary information

### 2 Supplementary table 1: sample overview different reproductive cycles split in main periods (5 periods).

| SB nr | Year | Status | Anestrus | Estrus | PostEstrus | CL dormancy | Active luteal | PostEndCycle | Total duration luteal phase | Date of peak oestrogen | First baseline P4 at end of cycle |
| --- | --- | --- | --- | --- | --- | --- | --- | --- | --- | --- | --- |
| 2014 | NB | Days from end-of-cycle | D-220/D-139 | D-140/D-139 | D-138/D-132 | D-131/D-43 | D-42/D0 | D1/D027 | 139 (+19 = 158) | 12 April 2014 | D19 |
|  |  | Days from peak estragen | D-81/D-11 | D-10/D0 | D-07/D01 | D-06/D008 | D-05/D0139 | D04/D0166 |  |  |  |
|  |  | Observed period (days) | 71 | 11 | 7 | 89 | 43 | 27 |  |  |  |
|  |  | Nr records estragens | 49 | 6 | 2 | 74 | 26 | 14 |  |  |  |
|  |  | Nr records progesterone | 21 | 6 | 2 | 74 | 26 | 14 |  |  |  |
|  |  | Nr records PGFM | 0 | 0 | 0 | 20 | 26 | 14 |  |  |  |
|  |  | Nr records CP | 0 | 0 | 1 | 72 | 26 | 13 |  |  |  |
|  |  | Nr records body weight | 68 | 2 | 4 | 86 | 24 | 5 |  |  |  |
|  |  | Nr records faecal output | 69 | 1 | 4 | 87 | 36 | 12 |  |  |  |
|  |  | 2015 | NB | Days from end-of-cycle | D-220/-58 | D-150/D-148 | D-147/D-141 | D-140/D-43 |  |  |  |
| Days from peak estragen | D-72/-12 |  |  | D-11/D00 | D-01/D07 | D-08/D005 | D-06/D0148 | D176 |  |  |  |
| Observed period (days) | 61 |  |  | 12 | 7 | 98 | 43 | 28 |  |  |  |
| Nr records estragens | 50 |  |  | 10 | 3 | 86 | 25 | 1 |  |  |  |
| Nr records progesterone | 50 |  |  | 10 | 3 | 86 | 24 | 1 |  |  |  |
| Nr records PGFM | 0 |  |  | 7 | 3 | 44 | 24 | 0 |  |  |  |
| Nr records CP | 32 |  |  | 10 | 3 | 82 | 24 | 1 |  |  |  |
| Nr records body weight | 59 |  |  | 6 | 2 | 98 | 27 | 3 |  |  |  |
| Nr records faecal output | 59 |  |  | 10 | 6 | 97 | 41 | 28 |  |  |  |
| 2016 | NB |  |  | Days from end-of-cycle | D-220/D-155 | D-154/D-145 | D-144/D-138 | D-137/D-43 | D-42/D0 | D1/D030 | 145 |
|  |  | Days from peak estragen | D-75/D-10 | D-9/D00 | D-01/D07 | D-08/D002 | D-03/D0145 | D146/D0175 |  |  |  |
|  |  | Observed period (days) | 66 | 10 | 7 | 95 | 43 | 30 |  |  |  |
|  |  | Nr records estragens | 58 | 7 | 1 | 75 | 35 | 5 |  |  |  |
|  |  | Nr records progesterone | 0 | 0 | 0 | 31 | 34 | 0 |  |  |  |
|  |  | Nr records PGFM | 2 | 7 | 1 | 23 | 34 | 5 |  |  |  |
|  |  | Nr records CP | 58 | 7 | 1 | 79 | 35 | 5 |  |  |  |
|  |  | Nr records body weight | 62 | 1 | 0 | 89 | 17 | 6 |  |  |  |
|  |  | Nr records faecal output | 66 | 9 | 5 | 93 | 42 | 30 |  |  |  |
|  |  | S8169 | NB | Days from end-of-cycle | D-220/D-179 | D-178/D-169 | D-168/D-162 | D-161/D-43 | D-42/D0 | D1/D014 |  |
| Days from peak estragen | D-51/D-10 |  |  | D-9/D00 | D-01/D07 | D-08/D0126 | D-01/D0169 | D170/D0183 |  |  |  |
| Observed period (days) | 62 |  |  | 10 | 7 | 119 | 43 | 14 |  |  |  |
| Nr records estragens | 39 |  |  | 8 | 3 | 102 | 39 | 6 |  |  |  |
| Nr records progesterone | 39 |  |  | 8 | 3 | 101 | 39 | 6 |  |  |  |
| Nr records PGFM | 6 |  |  | 8 | 3 | 38 | 39 | 6 |  |  |  |
| Nr records CP | 16 |  |  | 7 | 2 | 102 | 39 | 6 |  |  |  |
| Nr records body weight | 61 |  |  | 2 | 2 | 117 | 18 | 2 |  |  |  |
| Nr records faecal output | 61 |  |  | 7 | 6 | 119 | 42 | 18 |  |  |  |
| 2018 | PP |  |  | Days from end-of-cycle | D-220/D-163 | D-162/D-152 | D-151/D-145 | D-144/D-43 | D-42/D0 | D1/D024 | 152 (+16 = 168) |
|  |  | Days from peak estragen | D-48/D-11 | D-10/D00 | D-01/D07 | D-08/D0019 | D-01/D0132 | D134/D0176 |  |  |  |
|  |  | Observed period (days) | 58 | 11 | 7 | 102 | 43 | 24 |  |  |  |
|  |  | Nr records estragens | 29 | 7 | 4 | 81 | 16 | 5 |  |  |  |
|  |  | Nr records progesterone | 29 | 7 | 4 | 81 | 16 | 5 |  |  |  |
|  |  | Nr records PGFM | 0 | 0 | 0 | 31 | 16 | 5 |  |  |  |
|  |  | Nr records CP | 17 | 7 | 4 | 78 | 16 | 5 |  |  |  |
|  |  | Nr records body weight | 46 | 3 | 4 | 102 | 15 | 9 |  |  |  |
|  |  | Nr records faecal output | 55 | 9 | 7 | 102 | 38 | 22 |  |  |  |
|  |  | 2019 | NB | Days from end-of-cycle | D-220/D-162 | D-161/D-153 | D-152/D-146 | D-145/D-43 | D-42/D0 | D1/D030 |  |
| Days from peak estragen | D-67/D-9 |  |  | D-9/D00 | D-01/D07 | D-08/D0019 | D-01/D0153 | D134/D0183 |  |  |  |
| Observed period (days) | 59 |  |  | 9 | 7 | 101 | 43 | 30 |  |  |  |
| Nr records estragens | 52 |  |  | 9 | 7 | 97 | 39 | 17 |  |  |  |
| Nr records progesterone | 52 |  |  | 9 | 7 | 97 | 39 | 17 |  |  |  |
| Nr records PGFM | 0 |  |  | 0 | 0 | 13 | 39 | 17 |  |  |  |
| Nr records CP | 0 |  |  | 0 | 3 | 97 | 39 | 17 |  |  |  |
| Nr records body weight | 55 |  |  | 4 | 2 | 79 | 0 | 0 |  |  |  |
| Nr records faecal output | 59 |  |  | 9 | 7 | 102 | 43 | 26 |  |  |  |
| 2016 | NB |  |  | Days from end-of-cycle | D-220/D-139 | D-138/D-124 | D-122/D-117 | D-116/D-43 | D-42/D0 | D1/D030 | 124 (+18 = 142) |
|  |  | Days from peak estragen | D-96/D-15 | D-14/D00 | D-01/D07 | D-08/D001 | D-06/D0124 | D125/D0154 |  |  |  |
|  |  | Observed period (days) | 82 | 15 | 7 | 74 | 43 | 30 |  |  |  |
|  |  | Nr records estragens | 3 | 6 | 4 | 28 | 41 | 24 |  |  |  |
|  |  | Nr records progesterone | 3 | 6 | 5 | 72 | 41 | 24 |  |  |  |
|  |  | Nr records PGFM | 0 | 2 | 5 | 15 | 40 | 25 |  |  |  |
|  |  | Nr records CP | 0 | 1 | 8 | 1 | 41 | 25 |  |  |  |
|  |  | Nr records body weight | 12 | 2 | 1 | 10 | 7 | 3 |  |  |  |
|  |  | Nr records faecal output | 14 | 15 | 7 | 74 | 42 | 30 |  |  |  |
|  |  | S8723 | P (twin cub) | Days from end-of-cycle | D-220/D-149 | D-148/D-135 | D-134/D-128 | D-127/D-43 | D-42/D0 | D1/D030 |  |
| Days from peak estragen | D-85/D-14 |  |  | D-13/D00 | D-01/D07 | D-08/D002 | D-06/D0135 | D136/D0165 |  |  |  |
| Observed period (days) | 72 |  |  | 14 | 7 | 85 | 43 | 30 |  |  |  |
| Nr records estragens | 14 |  |  | 14 | 7 | 85 | 40 | 4 |  |  |  |
| Nr records progesterone | 14 |  |  | 13 | 7 | 85 | 40 | 4 |  |  |  |
| Nr records PGFM | 0 |  |  | 0 | 0 | 17 | 40 | 4 |  |  |  |
| Nr records CP | 10 |  |  | 14 | 7 | 85 | 40 | 3 |  |  |  |
| Nr records body weight | 11 |  |  | 2 | 1 | 12 | 5 | 4 |  |  |  |
| Nr records faecal output | 12 |  |  | 14 | 7 | 85 | 40 | 3 |  |  |  |
| 2020 | NB |  |  | Days from end-of-cycle | D-220/D-152 | D-150/D-96 | D-95/D-89 | D-88/D-43 | D-42/D0 | D1/D025 | 96 (+1 = 97) |
|  |  | Days from peak estragen | D-124/D-14 | D-13/D00 | D-01/D07 | D-08/D003 | D-05/D096 | D097/D0121 |  |  |  |
|  |  | Observed period (days) | 111 | 14 | 7 | 46 | 42 | 29 |  |  |  |
|  |  | Nr records estragens | 5 | 12 | 7 | 46 | 42 | 29 |  |  |  |
|  |  | Nr records progesterone | 5 | 12 | 7 | 46 | 42 | 29 |  |  |  |
|  |  | Nr records PGFM | 0 | 0 | 0 | 0 | 32 | 29 |  |  |  |
|  |  | Nr records CP | 5 | 12 | 7 | 46 | 42 | 29 |  |  |  |
|  |  | Nr records body weight | 16 | 2 | 1 | 7 | 6 | 4 |  |  |  |
|  |  | Nr records faecal output | 59 | 6 | 3 | 26 | 36 | 17 |  |  |  |
|  |  | 2015 | PP | Days from end-of-cycle | D-220/D-143 | D-142/D-132 | D-131/D-125 | D-124/D-43 | D-42/D0 | D1/D030 |  |
| Days from peak estragen | D-88/D-11 |  |  | D-10/D00 | D-01/D07 | D-08/D009 | D-06/D0132 | D133/D0162 |  |  |  |
| Observed period (days) | 78 |  |  | 11 | 7 | 82 | 43 | 30 |  |  |  |
| Nr records estragens | 51 |  |  | 8 | 4 | 11 | 9 | 6 |  |  |  |
| Nr records progesterone | 4 |  |  | 7 | 4 | 21 | 9 | 6 |  |  |  |
| Nr records PGFM | 0 |  |  | 7 | 4 | 21 | 9 | 6 |  |  |  |
| Nr records CP | 4 |  |  | 7 | 4 | 21 | 9 | 6 |  |  |  |
| Nr records body weight | 0 |  |  | 0 | 0 | 8 | 1 | 2 |  |  |  |
| Nr records faecal output | 0 |  |  | 0 | 0 | 78 | 41 | 23 |  |  |  |
| 2016 | P (single cub) |  |  | Days from end-of-cycle | D-129/D-127 | D-126/D-114 | D-113/D-107 | D-106/D-43 | D-42/D0 | D1/D012 | 114 |
|  |  | Days from peak estragen | D-15/D-13 | D-12/D00 | D-01/D07 | D-08/D001 | D-07/D0114 | D133/D0127 |  |  |  |
|  |  | Observed period (days) | 3 | 13 | 7 | 64 | 43 | 12 |  |  |  |
|  |  | Nr records estragens | 0 | 8 | 7 | 57 | 41 | 5 |  |  |  |
|  |  | Nr records progesterone | 0 | 8 | 7 | 57 | 41 | 5 |  |  |  |
|  |  | Nr records PGFM | 0 | 8 | 7 | 24 | 41 | 5 |  |  |  |
|  |  | Nr records CP | 0 | 8 | 7 | 57 | 41 | 5 |  |  |  |
|  |  | Nr records body weight | 0 | 0 | 0 | 0 | 0 | 0 |  |  |  |
|  |  | Nr records faecal output | 3 | 11 | 4 | 56 | 40 | 0 |  |  |  |
|  |  | 2018 | NB | Days from end-of-cycle | D-220/D-133 | D-132/D-122 | D-121/D-115 | D-114/D-43 | D-42/D0 | D1/D030 |  |
| Days from peak estragen | D-98/D-13 |  |  | D-10/D00 | D-01/D07 | D-08/D007 | D-06/D0122 | D123/D0152 |  |  |  |
| Observed period (days) | 88 |  |  | 11 | 7 | 72 | 43 | 30 |  |  |  |
| Nr records estragens | 73 |  |  | 11 | 7 | 69 | 39 | 29 |  |  |  |
| Nr records progesterone | 73 |  |  | 11 | 7 | 69 | 39 | 29 |  |  |  |
| Nr records PGFM | 0 |  |  | 0 | 0 | 9 | 39 | 29 |  |  |  |
| Nr records CP | 0 |  |  | 11 | 7 | 69 | 33 | 0 |  |  |  |
| Nr records body weight | 0 |  |  | 0 | 0 | 0 | 0 | 0 |  |  |  |
| Nr records faecal output | 26 |  |  | 8 | 6 | 64 | 39 | 27 |  |  |  |
| 2019 | P (twin cub) |  |  | Days from end-of-cycle | D-219/D-131 | D-133/D-111 | D-121/D-115 | D-114/D-43 | D-42/D0 | n.a. | 122 |
|  |  | Days from peak estragen | D-97/D-12 | D-11/D00 | D-01/D07 | D-08/D079 | D-06/D0122 | n.a. |  |  |  |
|  |  | Observed period (days) | 86 | 12 | 7 | 72 | 43 | 22 |  |  |  |
|  |  | Nr records estragens | 85 | 12 | 7 | 41 | 20 | n.a. |  |  |  |
|  |  | Nr records progesterone | 85 | 12 | 7 | 41 | 21 | n.a. |  |  |  |
|  |  | Nr records PGFM | 0 | 0 | 0 | 2 | 21 | n.a. |  |  |  |
|  |  | Nr records CP | 3 | 12 | 7 | 43 | 15 | n.a. |  |  |  |
|  |  | Nr records body weight | 0 | 0 | 0 | 0 | 0 | n.a. |  |  |  |
|  |  | Nr records faecal output | 86 | 12 | 2 | 72 | 39 | n.a. |  |  |  |
|  |  | 2017 | PP | Days from end-of-cycle | D-114/D-102 | D-100/D-91 | D-90/D-84 | D-83/D-43 | D-42/D0 | D1/D026 |  |
| Days from peak estragen | D-23/D-10 |  |  | D-9/D00 | D-01/D07 | D-08/D048 | D-06/D0121 | D062/D0117 |  |  |  |
| Observed period (days) | 14 |  |  | 10 | 7 | 41 | 43 | 28 |  |  |  |
| Nr records estragens | 2 |  |  | 2 | 2 | 1 | 21 | 0 |  |  |  |
| Nr records progesterone | 7 |  |  | 0 | 0 | 1 | 21 | 0 |  |  |  |
| Nr records PGFM | 2 |  |  | 2 | 2 | 1 | 21 | 0 |  |  |  |
| Nr records CP | 2 |  |  | 2 | 6 | 21 | 21 | 17 |  |  |  |
| Nr records body weight | 11 |  |  | 4 | 7 | 39 | 20 | 21 |  |  |  |
| Nr records faecal output | 8 |  |  | 5 | 7 | 41 | 29 | 21 |  |  |  |
| 2018 | PP |  |  | Days from end-of-cycle | D-220/-114 | D-113/D-102 | D-101/D-95 | D-94/D-43 | D-42/D0 | D1/D030 | 102 (+42 = 144) |
|  |  | Days from peak estragen | D-118/D-12 | D-11/D00 | D-01/D07 | D-08/D059 | D-06/D0102 | D103/D0132 |  |  |  |
|  |  | Observed period (days) | 107 | 12 | 7 | 55 | 37 | 22 |  |  |  |
|  |  | Nr records estragens | 70 | 11 | 5 | 37 | 22 | 21 |  |  |  |
|  |  | Nr records progesterone | 70 | 11 | 6 | 38 | 22 | 21 |  |  |  |
|  |  | Nr records PGFM | 4 | 4 | 11 | 10 | 22 | 21 |  |  |  |
|  |  | Nr records CP | 0 | 6 | 5 | 37 | 17 | 19 |  |  |  |
|  |  | Nr records body weight | 104 | 8 | 9 | 52 | 29 | 28 |  |  |  |
|  |  | Nr records faecal output | 105 | 12 | 4 | 52 | 29 | 28 |  |  |  |
|  |  | 2019 | PP | Days from end-of-cycle | D-220/D-124 | D-123/D-115 | D-114/D-108 | D-107/D-43 | D-42/D0 | D1/D030 |  |
| Days from peak estragen | D-105/D-9 |  |  | D-9/D00 | D-01/D07 | D-08/D072 | D-07/D0115 | D116/D0145 |  |  |  |
| Observed period (days) | 97 |  |  | 9 | 7 | 55 | 51 | 21 |  |  |  |
| Nr records estragens | 88 |  |  | 7 | 5 | 51 | 21 | 17 |  |  |  |
| Nr records progesterone | 88 |  |  | 7 | 5 | 51 | 27 | 17 |  |  |  |
| Nr records PGFM | 0 |  |  | 0 | 0 | 16 | 27 | n.a. |  |  |  |
| Nr records CP | 9 |  |  | 8 | 7 | 51 | 25 | n.a. |  |  |  |
| Nr records body weight | 98 |  |  | 11 | 5 | 50 | 37 | n.a. |  |  |  |
| Nr records faecal output | 104 |  |  | 11 | 5 | 55 | 39 | n.a. |  |  |  |
| 2020 | P (single cub) |  |  | Days from end-of-cycle | D-220/D-180 | D-158/D-148 | D-147/D-141 | D-140/D-43 | D-42/D0 | D1/D019 | 105 |
|  |  | Days from peak estragen | D-72/-12 | D-11/D00 | D-01/D07 | D-08/D005 | D-06/D0148 | D176/D0166 |  |  |  |
|  |  | Observed period (days) | 61 | 12 | 7 | 98 | 43 | 19 |  |  |  |
|  |  | Nr records estragens | 60 | 11 | 6 | 84 | 35 | 11 |  |  |  |
|  |  | Nr records progesterone | 60 | 11 | 6 | 84 | 35 | 11 |  |  |  |
|  |  | Nr records PGFM | 0 | 0 | 0 | 16 | 35 | 11 |  |  |  |
|  |  | Nr records CP | 6 | 11 | 6 | 84 | 35 | 11 |  |  |  |
|  |  | Nr records body weight | 5 | 1 | 1 | 9 | 0 | 0 |  |  |  |
|  |  | Nr records faecal output | 61 | 12 | 6 | 96 | 42 | 19 |  |  |  |
|  |  | S8684 | P (twin cub) | Days from end-of-cycle | D-220/D-172 | D-171/D-156 | D-155/D-149 | D-148/D-43 | D-42/D0 | D1/D030 |  |
| Days from peak estragen | D-64/D-16 |  |  | D-15/D00 | D-01/D07 | D-08/D013 | D-01/D0136 | D137/D0186 |  |  |  |
| Observed period (days) | 49 |  |  | 16 | 7 | 106 | 43 | 30 |  |  |  |
| Nr records estragens | 7 |  |  | 16 | 7 | 66 | 26 | 20 |  |  |  |
| Nr records progesterone | 7 |  |  | 16 | 7 | 68 | 26 | 20 |  |  |  |
| Nr records PGFM | 0 |  |  | 0 | 0 | 61 | 26 | 20 |  |  |  |
| Nr records CP | 0 |  |  | 0 | 1 | 61 | 26 | 20 |  |  |  |
| Nr records body weight | 11 |  |  | 6 | 3 | 23 | 13 | 5 |  |  |  |
| Nr records faecal output | 11 |  |  | 6 | 3 | 23 | 13 | 5 |  |  |  |
| S8941 | PP |  |  | Days from end-of-cycle | D-220/D-172 | D-171/D-156 | D-155/D-149 | D-148/D-43 | D-42/D0 | D1/D030 | 156 (+4 = 160) |
|  |  | Days from peak estragen | D-64/D-16 | D-15/D00 | D-01/D07 | D-08/D013 | D-01/D0136 | D137/D0186 |  |  |  |
|  |  | Observed period (days) | 49 | 16 | 7 | 106 | 43 | 30 |  |  |  |
|  |  | Nr records estragens | 7 | 16 | 7 | 66 | 26 | 20 |  |  |  |
|  |  | Nr records progesterone | 7 | 16 | 7 | 68 | 26 | 20 |  |  |  |
|  |  | Nr records PGFM | 0 | 0 | 0 | 61 | 26 | 20 |  |  |  |
|  |  | Nr records CP | 0 | 0 | 1 | 61 | 26 | 20 |  |  |  |
|  |  | Nr records body weight | 11 | 6 | 3 | 23 | 13 | 5 |  |  |  |
|  |  | Nr records faecal output | 11 | 6 | 3 | 23 | 13 | 5 |  |  |  |

4 Supplementary table 2: sample overview different reproductive cycles split in subperiods for CLD and AL.

| SR nr | Year | Status | Corpus Luteum Dormancy Phase |  |  |  |  |  |  |  |  |  | Active Luteal Phase |  |  |  |  |  |  |
| --- | --- | --- | --- | --- | --- | --- | --- | --- | --- | --- | --- | --- | --- | --- | --- | --- | --- | --- | --- |
|  |  |  | I |  | II |  | III |  | 1 |  | 2 |  | 3 |  | 4 |  | 5 |  | 6 |
| 2014 | NB |  | Days from end-of-cycle | D-131/D-97 | D-96/D-60 | D-59/D-43 | D-42/D-36 | D-35/29 | D-28/D-22 | D-21/D-15 | D-14/D-8 | D-7/D0 |  |  |  |  |  |  |  |
|  |  |  | Days from peak estragen | D6/D42 | D43/D79 | D80/D96 | D97/D103 | D104/D110 | D111/D117 | D118/D124 | D125/D131 | D132/D139 |  |  |  |  |  |  |  |
|  |  |  | Observed period (days) | 35 | 37 | 17 | 7 | 7 | 7 | 7 | 7 | 7 | 8 |  |  |  |  |  |  |
|  |  |  | Nr records estragens | 27 | 31 | 16 | 3 | 1 | 4 | 3 | 7 | 7 | 8 |  |  |  |  |  |  |
|  |  |  | Nr records progesterone | 27 | 31 | 16 | 3 | 1 | 4 | 3 | 7 | 7 | 8 |  |  |  |  |  |  |
|  |  |  | Nr records PGFM | 0 | 4 | 16 | 3 | 1 | 4 | 3 | 7 | 7 | 8 |  |  |  |  |  |  |
|  |  |  | Nr records CP | 26 | 31 | 15 | 3 | 1 | 4 | 3 | 7 | 7 | 8 |  |  |  |  |  |  |
|  |  |  | Nr records body weight | 34 | 36 | 16 | 7 | 7 | 3 | 2 | 1 | 4 | 4 |  |  |  |  |  |  |
|  |  |  | Nr records faecal output | 35 | 36 | 16 | 7 | 7 | 3 | 2 | 1 | 4 | 4 |  |  |  |  |  |  |
|  |  |  | Nr records facial output | 35 | 36 | 16 | 7 | 7 | 3 | 2 | 1 | 4 | 4 |  |  |  |  |  |  |
| 2015 | NB |  | Days from end-of-cycle | D-140/D-83 | D-82/D-63 | D-62/D-43 | D-42/D-36 | D-35/29 | D-28/D-22 | D-21/D-15 | D-14/D-8 | D-7/D0 |  |  |  |  |  |  |  |
|  |  |  | Days from peak estragen | D6/D65 | D66/D85 | D86/D105 | D106/D112 | D113/D119 | D120/D126 | D127/D133 | D134/D140 | D141/D148 |  |  |  |  |  |  |  |
|  |  |  | Observed period (days) | 58 | 20 | 20 | 7 | 7 | 7 | 7 | 7 | 7 | 8 |  |  |  |  |  |  |
|  |  |  | Nr records estragens | 53 | 18 | 15 | 7 | 5 | 5 | 3 | 4 | 1 | 1 |  |  |  |  |  |  |
|  |  |  | Nr records progesterone | 53 | 18 | 15 | 7 | 5 | 5 | 4 | 3 | 4 | 1 |  |  |  |  |  |  |
|  |  |  | Nr records PGFM | 11 | 18 | 15 | 7 | 5 | 5 | 4 | 3 | 4 | 1 |  |  |  |  |  |  |
|  |  |  | Nr records CP | 51 | 16 | 15 | 7 | 5 | 5 | 4 | 3 | 4 | 1 |  |  |  |  |  |  |
|  |  |  | Nr records body weight | 58 | 20 | 20 | 7 | 7 | 5 | 4 | 2 | 2 | 2 |  |  |  |  |  |  |
|  |  |  | Nr records faecal output | 58 | 20 | 20 | 7 | 7 | 5 | 4 | 2 | 2 | 2 |  |  |  |  |  |  |
|  |  |  | Nr records facial output | 58 | 20 | 20 | 7 | 7 | 5 | 4 | 2 | 2 | 2 |  |  |  |  |  |  |
| 2016 | NB |  | Days from end-of-cycle | D-137/D-75 | D-74/D-61 | D-60/D-43 | D-42/D-36 | D-35/29 | D-28/D-22 | D-21/D-15 | D-14/D-8 | D-7/D0 |  |  |  |  |  |  |  |
|  |  |  | Days from peak estragen | D6/D70 | D71/D84 | D85/D102 | D103/D109 | D110/D116 | D117/D123 | D124/D130 | D131/D137 | D138/D145 |  |  |  |  |  |  |  |
|  |  |  | Observed period (days) | 63 | 14 | 18 | 7 | 7 | 7 | 7 | 7 | 7 | 8 |  |  |  |  |  |  |
|  |  |  | Nr records estragens | 48 | 11 | 16 | 4 | 6 | 6 | 7 | 7 | 7 | 5 |  |  |  |  |  |  |
|  |  |  | Nr records progesterone | 7 | 10 | 14 | 4 | 5 | 6 | 7 | 7 | 5 | 7 |  |  |  |  |  |  |
|  |  |  | Nr records PGFM | 4 | 8 | 11 | 4 | 5 | 6 | 7 | 7 | 5 | 7 |  |  |  |  |  |  |
|  |  |  | Nr records CP | 48 | 13 | 18 | 4 | 6 | 8 | 7 | 7 | 5 | 7 |  |  |  |  |  |  |
|  |  |  | Nr records body weight | 58 | 14 | 17 | 7 | 7 | 3 | 0 | 0 | 0 | 0 |  |  |  |  |  |  |
|  |  |  | Nr records faecal output | 62 | 13 | 18 | 7 | 7 | 7 | 7 | 7 | 7 | 8 |  |  |  |  |  |  |
|  |  |  | Nr records facial output | 62 | 13 | 18 | 7 | 7 | 7 | 7 | 7 | 7 | 8 |  |  |  |  |  |  |
| 2017 | NB |  | Days from end-of-cycle | D-162/D-83 | D-82/D-65 | D-64/D-43 | D-42/D-36 | D-35/29 | D-28/D-22 | D-21/D-15 | D-14/D-8 | D-7/D0 |  |  |  |  |  |  |  |
|  |  |  | Days from peak estragen | D6/D86 | D87/D104 | D105/D126 | D127/D133 | D134/D140 | D141/D147 | D148/D154 | D155/D161 | D162/D169 |  |  |  |  |  |  |  |
|  |  |  | Observed period (days) | 79 | 18 | 22 | 7 | 7 | 7 | 7 | 7 | 7 | 8 |  |  |  |  |  |  |
|  |  |  | Nr records estragens | 71 | 14 | 17 | 6 | 6 | 7 | 7 | 7 | 7 | 6 |  |  |  |  |  |  |
|  |  |  | Nr records progesterone | 71 | 13 | 17 | 6 | 6 | 7 | 7 | 7 | 7 | 6 |  |  |  |  |  |  |
|  |  |  | Nr records PGFM | 7 | 14 | 17 | 6 | 6 | 7 | 7 | 7 | 7 | 6 |  |  |  |  |  |  |
|  |  |  | Nr records CP | 71 | 14 | 17 | 6 | 6 | 7 | 7 | 7 | 7 | 6 |  |  |  |  |  |  |
|  |  |  | Nr records body weight | 79 | 18 | 22 | 7 | 7 | 7 | 7 | 7 | 7 | 8 |  |  |  |  |  |  |
|  |  |  | Nr records faecal output | 79 | 18 | 22 | 7 | 7 | 7 | 7 | 7 | 7 | 8 |  |  |  |  |  |  |
|  |  |  | Nr records facial output | 79 | 18 | 22 | 7 | 7 | 7 | 7 | 7 | 7 | 8 |  |  |  |  |  |  |
| 2018 | PP |  | Days from end-of-cycle | D-144/D-89 | D-88/D-70 | D-69/D-43 | D-42/D-36 | D-35/29 | D-28/D-22 | D-21/D-15 | D-14/D-8 | D-7/D0 |  |  |  |  |  |  |  |
|  |  |  | Days from peak estragen | D6/D93 | D94/D102 | D103/D109 | D110/D116 | D117/D123 | D124/D130 | D131/D137 | D138/D144 | D145/D152 |  |  |  |  |  |  |  |
|  |  |  | Observed period (days) | 56 | 19 | 27 | 7 | 7 | 7 | 7 | 7 | 7 | 8 |  |  |  |  |  |  |
|  |  |  | Nr records estragens | 48 | 13 | 20 | 4 | 2 | 4 | 4 | 4 | 1 | 1 |  |  |  |  |  |  |
|  |  |  | Nr records progesterone | 48 | 13 | 20 | 4 | 2 | 4 | 4 | 4 | 1 | 1 |  |  |  |  |  |  |
|  |  |  | Nr records PGFM | 0 | 11 | 20 | 4 | 2 | 4 | 4 | 4 | 1 | 1 |  |  |  |  |  |  |
|  |  |  | Nr records CP | 45 | 13 | 20 | 4 | 2 | 4 | 4 | 4 | 1 | 1 |  |  |  |  |  |  |
|  |  |  | Nr records body weight | 56 | 19 | 27 | 7 | 7 | 7 | 7 | 7 | 7 | 8 |  |  |  |  |  |  |
|  |  |  | Nr records faecal output | 56 | 19 | 27 | 7 | 7 | 7 | 7 | 7 | 7 | 8 |  |  |  |  |  |  |
|  |  |  | Nr records facial output | 56 | 19 | 27 | 7 | 7 | 7 | 7 | 7 | 7 | 8 |  |  |  |  |  |  |
| 2019 | NB |  | Days from end-of-cycle | D-145/D-97 | D-96/D-66 | D-65/D-43 | D-42/D-36 | D-35/29 | D-28/D-22 | D-21/D-15 | D-14/D-8 | D-7/D0 |  |  |  |  |  |  |  |
|  |  |  | Days from peak estragen | D6/D102 | D103/D107 | D108/D110 | D111/D117 | D118/D124 | D125/D131 | D132/D138 | D139/D145 | D146/D153 |  |  |  |  |  |  |  |
|  |  |  | Observed period (days) | 49 | 31 | 23 | 7 | 7 | 7 | 7 | 7 | 7 | 8 |  |  |  |  |  |  |
|  |  |  | Nr records estragens | 48 | 26 | 23 | 6 | 7 | 6 | 7 | 6 | 7 | 7 |  |  |  |  |  |  |
|  |  |  | Nr records progesterone | 48 | 26 | 23 | 6 | 7 | 6 | 7 | 6 | 7 | 7 |  |  |  |  |  |  |
|  |  |  | Nr records PGFM | 0 | 0 | 13 | 6 | 7 | 6 | 7 | 6 | 7 | 7 |  |  |  |  |  |  |
|  |  |  | Nr records CP | 48 | 26 | 23 | 6 | 7 | 6 | 7 | 6 | 7 | 7 |  |  |  |  |  |  |
|  |  |  | Nr records body weight | 45 | 31 | 3 | 0 | 0 | 0 | 0 | 0 | 0 | 0 |  |  |  |  |  |  |
|  |  |  | Nr records faecal output | 48 | 31 | 23 | 7 | 7 | 7 | 7 | 7 | 7 | 8 |  |  |  |  |  |  |
|  |  |  | Nr records facial output | 48 | 31 | 23 | 7 | 7 | 7 | 7 | 7 | 7 | 8 |  |  |  |  |  |  |
| 2020 | NB |  | Days from end-of-cycle | D-166/D-83 | D-82/D-62 | D-61/D-43 | D-42/D-36 | D-35/29 | D-28/D-22 | D-21/D-15 | D-14/D-8 | D-7/D0 |  |  |  |  |  |  |  |
|  |  |  | Days from peak estragen | D6/D94 | D95/D103 | D104/D112 | D113/D121 | D120/D128 | D129/D137 | D138/D146 | D147/D155 | D156/D164 |  |  |  |  |  |  |  |
|  |  |  | Observed period (days) | 34 | 21 | 19 | 7 | 7 | 7 | 7 | 7 | 7 | 8 |  |  |  |  |  |  |
|  |  |  | Nr records estragens | 35 | 8 | 2 | 18 | 7 | 7 | 7 | 7 | 7 | 6 |  |  |  |  |  |  |
|  |  |  | Nr records progesterone | 33 | 20 | 19 | 7 | 7 | 7 | 7 | 7 | 7 | 6 |  |  |  |  |  |  |
|  |  |  | Nr records PGFM | 15 | 0 | 0 | 6 | 7 | 7 | 7 | 7 | 7 | 6 |  |  |  |  |  |  |
|  |  |  | Nr records CP | 35 | 8 | 7 | 8 | 7 | 7 | 7 | 7 | 7 | 6 |  |  |  |  |  |  |
|  |  |  | Nr records body weight | 5 | 3 | 2 | 1 | 1 | 1 | 1 | 1 | 1 | 2 |  |  |  |  |  |  |
|  |  |  | Nr records faecal output | 44 | 24 | 17 | 7 | 7 | 7 | 7 | 7 | 7 | 8 |  |  |  |  |  |  |
|  |  |  | Nr records facial output | 44 | 24 | 17 | 7 | 7 | 7 | 7 | 7 | 7 | 8 |  |  |  |  |  |  |
| 2021 | P (twin cub) |  | Days from end-of-cycle | D-127/D-84 | D-83/D-60 | D-59/D-43 | D-42/D-36 | D-35/29 | D-28/D-22 | D-21/D-15 | D-14/D-8 | D-7/D0 |  |  |  |  |  |  |  |
|  |  |  | Days from peak estragen | D6/D105 | D106/D113 | D114/D121 | D122/D129 | D130/D137 | D138/D145 | D146/D153 | D154/D161 | D162/D169 |  |  |  |  |  |  |  |
|  |  |  | Observed period (days) | 44 | 24 | 17 | 7 | 7 | 7 | 7 | 7 | 7 | 8 |  |  |  |  |  |  |
|  |  |  | Nr records estragens | 44 | 24 | 17 | 7 | 7 | 7 | 7 | 7 | 7 | 5 |  |  |  |  |  |  |
|  |  |  | Nr records progesterone | 44 | 24 | 17 | 7 | 7 | 7 | 7 | 7 | 7 | 5 |  |  |  |  |  |  |
|  |  |  | Nr records PGFM | 4 | 24 | 17 | 7 | 7 | 7 | 7 | 7 | 7 | 5 |  |  |  |  |  |  |
|  |  |  | Nr records CP | 44 | 24 | 17 | 7 | 7 | 7 | 7 | 7 | 7 | 5 |  |  |  |  |  |  |
|  |  |  | Nr records body weight | 6 | 3 | 3 | 1 | 1 | 1 | 1 | 1 | 1 | 1 |  |  |  |  |  |  |
|  |  |  | Nr records faecal output | 44 | 24 | 17 | 7 | 7 | 7 | 7 | 7 | 7 | 5 |  |  |  |  |  |  |
|  |  |  | Nr records facial output | 44 | 24 | 17 | 7 | 7 | 7 | 7 | 7 | 7 | 5 |  |  |  |  |  |  |
| 2022 | NB |  | Days from end-of-cycle | D-88/D-71 | D-70/D-58 | D-57/D-43 | D-42/D-36 | D-35/29 | D-28/D-22 | D-21/D-15 | D-14/D-8 | D-7/D0 |  |  |  |  |  |  |  |
|  |  |  | Days from peak estragen | D6/D125 | D126/D138 | D139/D151 | D150/D160 | D161/D171 | D172/D182 | D183/D193 | D194/D203 | D204/D213 |  |  |  |  |  |  |  |
|  |  |  | Observed period (days) | 35 | 13 | 15 | 7 | 7 | 7 | 7 | 7 | 7 | 8 |  |  |  |  |  |  |
|  |  |  | Nr records estragens | 38 | 13 | 15 | 7 | 7 | 7 | 7 | 7 | 7 | 7 |  |  |  |  |  |  |
|  |  |  | Nr records progesterone | 38 | 13 | 15 | 7 | 7 | 7 | 7 | 7 | 7 | 7 |  |  |  |  |  |  |
|  |  |  | Nr records PGFM | 0 | 0 | 0 | 4 | 7 | 7 | 7 | 7 | 7 | 7 |  |  |  |  |  |  |
|  |  |  | Nr records CP | 38 | 13 | 15 | 7 | 7 | 7 | 7 | 7 | 7 | 7 |  |  |  |  |  |  |
|  |  |  | Nr records body weight | 3 | 2 | 2 | 5 | 1 | 1 | 1 | 1 | 1 | 1 |  |  |  |  |  |  |
|  |  |  | Nr records faecal output | 31 | 7 | 8 | 5 | 5 | 5 | 5 | 5 | 5 | 7 |  |  |  |  |  |  |
|  |  |  | Nr records facial output | n.a. | n.a. | n.a. | n.a. | n.a. | n.a. | n.a. | n.a. | n.a. | n.a. |  |  |  |  |  |  |
| 2023 | PP |  | Days from end-of-cycle | n.a. | n.a. | n.a. | D-42/D-36 | D-35/29 | D-28/D-22 | D-21/D-15 | D-14/D-8 | D-7/D0 |  |  |  |  |  |  |  |
|  |  |  | Days from peak estragen | n.a. | n.a. | n.a. | D56/D66 | D67/D75 | D76/D84 | D85/D93 | D94/D101 | D102/D110 |  |  |  |  |  |  |  |
|  |  |  | Observed period (days) | n.a. | n.a. | n.a. | 096/096 | 097/103 | 104/110 | 111/117 | 118/124 | 125/132 |  |  |  |  |  |  |  |
|  |  |  | Nr records estragens | n.a. | n.a. | n.a. | 1 | 1 | 2 | 1 | 1 | 1 |  |  |  |  |  |  |  |
|  |  |  | Nr records progesterone | n.a. | n.a. | n.a. | 1 | 1 | 2 | 1 | 1 | 1 |  |  |  |  |  |  |  |
|  |  |  | Nr records PGFM | n.a. | n.a. | n.a. | 1 | 1 | 2 | 1 | 1 | 1 |  |  |  |  |  |  |  |
|  |  |  | Nr records CP | n.a. | n.a. | n.a. | 1 | 1 | 2 | 1 | 1 | 1 |  |  |  |  |  |  |  |
|  |  |  | Nr records body weight | n.a. | n.a. | n.a. | n.a. | n.a. | 1 | 1 | 2 | 1 |  |  |  |  |  |  |  |
|  |  |  | Nr records faecal output | n.a. | n.a. | n.a. | n.a. | n.a. | n.a. | n.a. | n.a. | n.a. | n.a. |  |  |  |  |  |  |
|  |  |  | Nr records facial output | n.a. | n.a. | n.a. | n.a. | n.a. | n.a. | n.a. | n.a. | n.a. | n.a. |  |  |  |  |  |  |
| 2024 | P (single cub) |  | Days from end-of-cycle | D-106/D-88 | D-85/D-61 | D-61/D-43 | D-42/D-36 | D-35/29 | D-28/D-22 | D-21/D-15 | D-14/D-8 | D-7/D0 |  |  |  |  |  |  |  |
|  |  |  | Days from peak estragen | D6/D104 | D105/D113 | D114/D121 | D122/D129 | D130/D137 | D138/D145 | D146/D153 | D154/D161 | D162/D169 |  |  |  |  |  |  |  |
|  |  |  | Observed period (days) | 27 | 19 | 18 | 7 | 7 | 7 | 7 | 7 | 7 | 8 |  |  |  |  |  |  |
|  |  |  | Nr records estragens | 25 | 17 | 15 | 7 | 7 | 7 | 7 | 7 | 7 | 6 |  |  |  |  |  |  |
|  |  |  | Nr records progesterone | 25 | 17 | 15 | 7 | 7 | 7 | 7 | 7 | 7 | 6 |  |  |  |  |  |  |
|  |  |  | Nr records PGFM | 17 | 0 | 7 | 7 | 7 | 7 | 7 | 7 | 7 | 6 |  |  |  |  |  |  |
|  |  |  | Nr records CP | 25 | 17 | 15 | 7 | 7 | 7 | 7 | 7 | 7 | 6 |  |  |  |  |  |  |
|  |  |  | Nr records body weight | n.a. | n.a. | n.a. | n.a. | n.a. | n.a. | n.a. | n.a. | n.a. | n.a. |  |  |  |  |  |  |
|  |  |  | Nr records faecal output | 24 | 17 | 15 | 7 | 7 | 7 | 7 | 7 | 7 | 6 |  |  |  |  |  |  |
|  |  |  | Nr records facial output | n.a. | n.a. | n.a. | n.a. | n.a. | n.a. | n.a. | n.a. | n.a. | n.a. |  |  |  |  |  |  |
| 2025 | NB |  | Days from end-of-cycle | D-114/D-89 | D-88/D-62 | D-61/D-43 | D-42/D-36 | D-35/29 | D-28/D-22 | D-21/D-15 | D-14/D-8 | D-7/D0 |  |  |  |  |  |  |  |
|  |  |  | Days from peak estragen | D6/D131 | D132/D140 | D141/D149 | D150/D158 | D159/D167 | D168/D176 | D177/D185 | D186/D194 | D195/D203 |  |  |  |  |  |  |  |
|  |  |  | Observed period (days) | 26 | 27 | 19 | 7 | 7 | 7 | 7 | 7 | 7 | 8 |  |  |  |  |  |  |
|  |  |  | Nr records estragens | 26 | 24 | 19 | 6 | 6 | 6 | 7 | 7 | 5 | 8 |  |  |  |  |  |  |
|  |  |  | Nr records progesterone | 26 | 24 | 19 | 6 | 6 | 6 | 7 | 7 | 5 | 8 |  |  |  |  |  |  |
|  |  |  | Nr records PGFM | 0 | 0 | 9 | 6 | 6 | 7 | 7 | 7 | 5 | 8 |  |  |  |  |  |  |
|  |  |  | Nr records CP | 26 | 24 | 19 | 6 | 6 | 6 | 7 | 7 | 5 | 8 |  |  |  |  |  |  |
|  |  |  | Nr records body weight | n.a. | n.a. | n.a. | n.a. | n.a. | n.a. | n.a. | n.a. | n.a. | n.a. |  |  |  |  |  |  |
|  |  |  | Nr records faecal output | n.a. | n.a. | n.a. | n.a. | n.a. | n.a. | n.a. | n.a. | n.a. | n.a. |  |  |  |  |  |  |
|  |  |  | Nr records facial output | n.a. | n.a. | n.a. | n.a. | n.a. | n.a. | n.a. | n.a. | n.a. | n.a. |  |  |  |  |  |  |
| 2026 | P (twin cub) |  | Days from end-of-cycle | D-114/D-74 | D-73/D-57 | D-56/D-43 | D-42/D-36 | D-35/29 | D-28/D-22 | D-21/D-15 | D-14/D-8 | D-7/D0 |  |  |  |  |  |  |  |
|  |  |  | Days from peak estragen | D6/D48 | D49/D65 | D66/D79 | D80/D86 | D87/D93 | D94/D100 | D101/D107 | D108/D114 | D115/D122 |  |  |  |  |  |  |  |
|  |  |  | Observed period (days) | 41 | 17 | 14 | 7 | 7 | 7 | 7 | 7 | 7 | 8 |  |  |  |  |  |  |
|  |  |  | Nr records estragens | n.a. | 27 | 9 | 7 | 7 | 7 | 7 | 7 | 7 | 5 |  |  |  |  |  |  |
|  |  |  | Nr records progesterone | n.a. | 27 | 9 | 5 | 0 | 7 | 6 | 0 | 2 | 6 |  |  |  |  |  |  |
|  |  |  | Nr records PGFM | n.a. | 27 | 9 | 5 | 0 | 7 | 6 | 0 | 2 | 6 |  |  |  |  |  |  |
|  |  |  | Nr records CP | n.a. | 27 | 9 | 5 | 0 | 7 | 6 | 0 | 2 | 6 |  |  |  |  |  |  |
|  |  |  | Nr records body weight | n.a. | n.a. | n.a. | n.a. | n.a. | n.a. | n.a. | n.a. | n.a. | n.a. |  |  |  |  |  |  |
|  |  |  | Nr records faecal output | n.a. | n.a. | n.a. | n.a. | n.a. | n.a. | n.a. | n.a. | n.a. | n.a. |  |  |  |  |  |  |
|  |  |  | Nr records facial output | n.a. | n.a. | n.a. | n.a. | n.a. | n.a. | n.a. | n.a. | n.a. | n.a. |  |  |  |  |  |  |
| 2027 | PP |  |  |  |  |  |  |  |  |  |  |  |  |  |  |  |  |  |  |

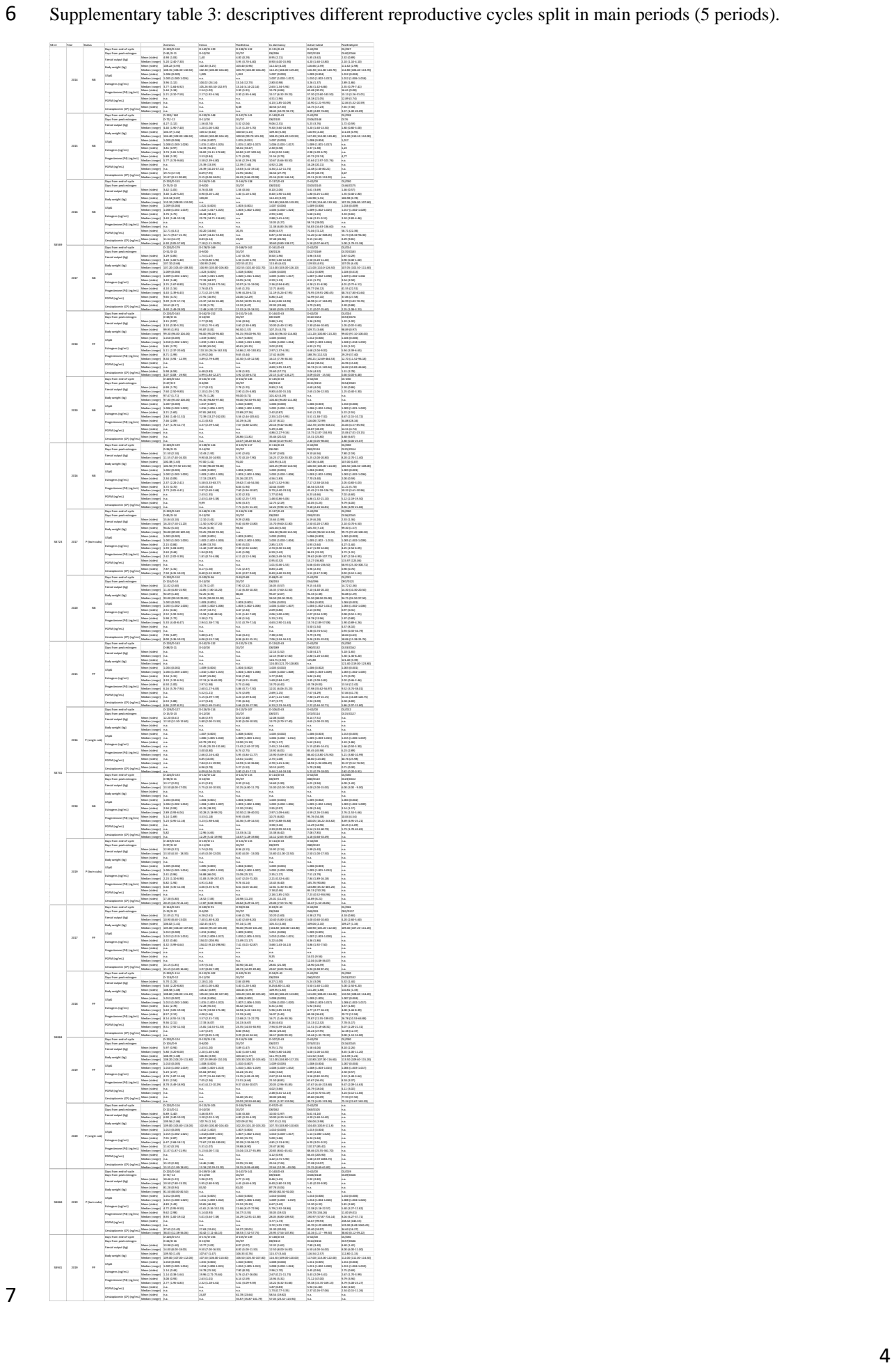

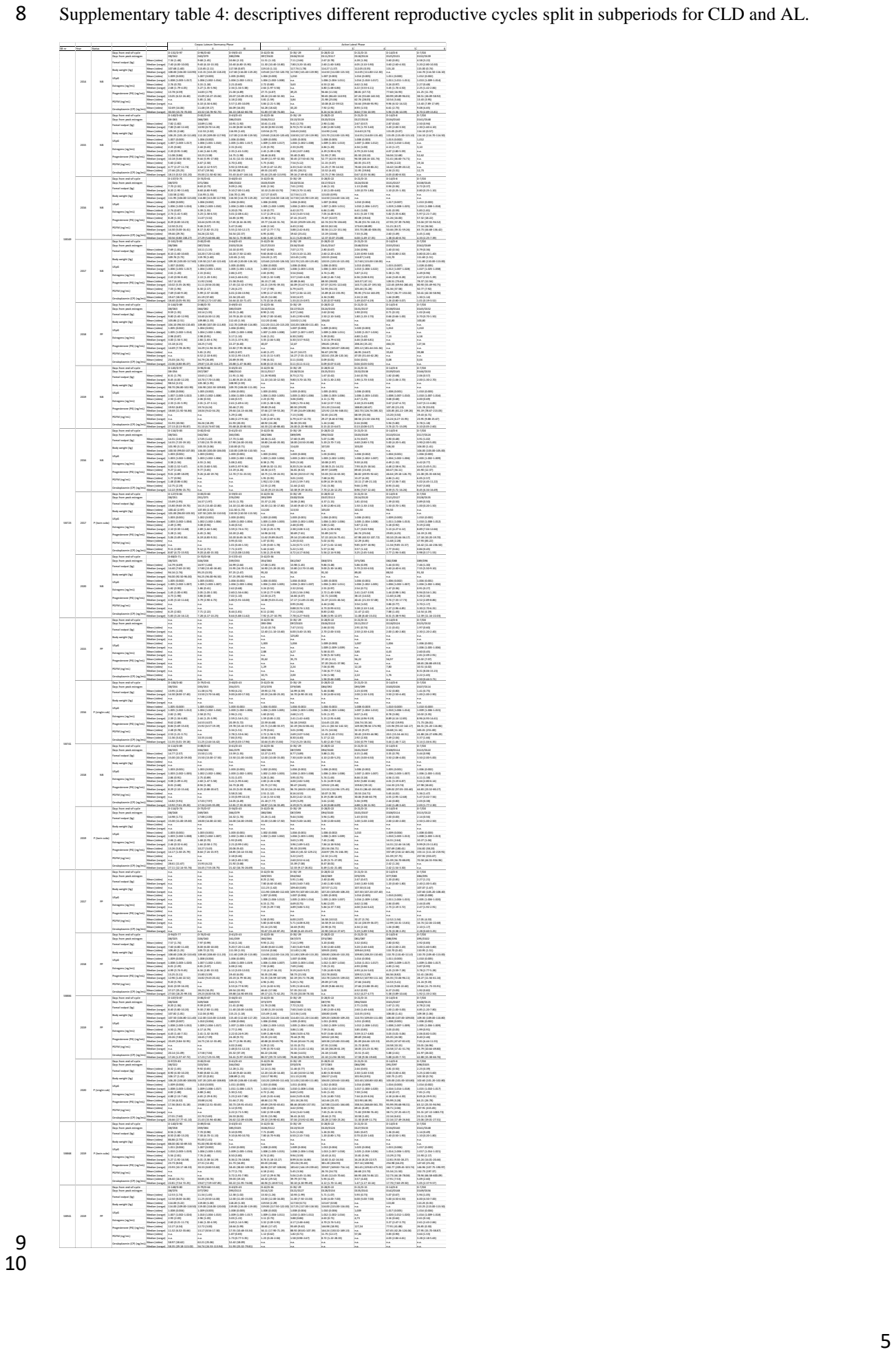

11 Supplementary table 5: additional data on the lenght of the different CLD phases per category of reproductive
12 cycles (pregnant – pseudopregnant – non-birth).

| Category |  | Average anestrus bodyweight | Average postestrus bodyweight | Dormancy I |  |  | Dormancy II |  |  | Dormancy III |  |  | Total CL dormancy |  |  |
| --- | --- | --- | --- | --- | --- | --- | --- | --- | --- | --- | --- | --- | --- | --- | --- |
|  |  |  |  | Start | Stop | Interval | Start | Stop | Interval | Start | Stop | Interval | Start | Stop | Interval |
| Pregnant | Days from end-of-cycle (-) |  |  | 106 | 80 |  | 79 | 61 |  | 60 | 43 |  | 106 | 43 |  |
|  | Days from peak estrogen (+) | 120 |  | 8 | 34 | 27 | 35 | 53 | 19 | 54 | 71 | 18 | 8 | 71 | 64 |
|  | Days from end-of-cycle (-) |  |  | 114 | 74 |  | 73 | 57 |  | 56 | 43 |  | 114 | 43 |  |
| HH2019 | Days from peak estrogen (+) | 120 |  | 8 | 48 | 41 | 49 | 65 | 17 | 66 | 79 | 14 | 8 | 79 | 72 |
| HuHu2017 | Days from end-of-cycle (-) |  |  | 127 | 84 |  | 83 | 60 |  | 59 | 43 |  | 127 | 43 |  |
|  | Days from peak estrogen (+) | 94,82 | 93,5 | 8 | 51 | 44 | 52 | 75 | 24 | 76 | 92 | 17 | 8 | 92 | 85 |
|  | Days from end-of-cycle (-) |  |  | 140 | 90 |  | 89 | 64 |  | 63 | 43 |  | 140 | 43 |  |
| MM2019 | Days from peak estrogen (+) | 81,28 | 81 | 8 | 58 | 51 | 59 | 84 | 26 | 85 | 105 | 21 | 8 | 105 | 98 |
| WW2020 | Days from end-of-cycle (-) |  |  | 97 | 83 |  | 82 | 62 |  | 61 | 43 |  | 97 | 43 |  |
|  | Days from peak estrogen (+) | 109,56 | 102,09 | 8 | 22 | 15 | 23 | 43 | 21 | 44 | 62 | 19 | 8 | 62 | 55 |
|  | Days from end-of-cycle (-) |  |  | 116,80 | 82,20 | 35,60 | 81,20 | 60,80 | 21,40 | 59,80 | 43,00 | 17,80 | 116,80 | 43,00 | 74,80 |
| Average | Days from end-of-cycle (-) |  |  | 17,02 | 5,85 | 14,45 | 5,85 | 2,59 | 3,65 | 2,59 | 0,00 | 2,59 | 17,02 | 0,00 | 17,02 |
| Std | Days from end-of-cycle (-) |  |  | 8,00 | 42,60 |  | 43,60 | 64,00 |  | 65,00 | 81,80 |  | 8,00 | 81,80 |  |
| Average | Days from peak estrogen (+) |  |  | 0,00 | 14,45 |  | 14,45 | 16,46 |  | 16,46 | 17,02 |  | 0,00 | 17,02 |  |
| Std | Days from peak estrogen (+) |  |  |  |  |  |  |  |  |  |  |  |  |  |  |
| Pseudopregnant |  |  |  |  |  |  |  |  |  |  |  |  |  |  |  |
| HH2015 | Days from end-of-cycle (-) |  |  |  |  |  |  |  |  |  |  |  | 124 | 43 |  |
|  | Days from peak estrogen (+) | 120 |  |  |  |  |  |  |  |  |  |  | 8 | 89 | 82 |
|  | Days from end-of-cycle (-) |  |  | 144 | 89 |  | 88 | 70 |  | 69 | 43 |  | 144 | 43 |  |
| TT2018 | Days from peak estrogen (+) | 99,95 | 94,5 | 8 | 63 | 55 | 64 | 82 | 19 | 83 | 109 | 27 | 8 | 109 | 101 |
| WW2017 | Days from end-of-cycle (-) |  |  |  |  |  |  |  |  |  |  |  | 83 | 43 |  |
|  | Days from peak estrogen (+) | 106,02 | 97,14 |  |  |  |  |  |  |  |  |  | 8 | 48 | 41 |
|  | Days from end-of-cycle (-) |  |  | 94 | 77 |  | 76 | 62 |  | 61 | 43 |  | 94 | 43 |  |
| WW2018 | Days from peak estrogen (+) | 108,58 | 104,45 | 8 | 25 | 18 | 26 | 40 | 15 | 41 | 59 | 19 | 8 | 59 | 52 |
| WW2019 | Days from end-of-cycle (-) |  |  | 107 | 87 |  | 86 | 67 |  | 66 | 43 |  | 107 | 43 |  |
|  | Days from peak estrogen (+) | 108,39 | 103,3 | 8 | 28 | 21 | 29 | 48 | 20 | 49 | 72 | 24 | 8 | 72 | 65 |
|  | Days from end-of-cycle (-) |  |  | 148 | 80 |  | 79 | 64 |  | 63 | 43 |  | 148 | 43 |  |
| JBB2019 | Days from peak estrogen (+) | 109,5 | 106,33 | 8 | 76 | 69 | 77 | 92 | 16 | 93 | 113 | 21 | 8 | 113 | 106 |
| Average | Days from end-of-cycle (-) |  |  | 123,25 | 83,25 | 40,75 | 82,25 | 65,75 | 17,50 | 64,75 | 43,00 | 22,75 | 115,20 | 43,00 | 73,00 |
|  | Days from end-of-cycle (-) |  |  | 26,85 | 5,68 | 25,22 | 5,68 | 3,50 | 2,38 | 3,50 | 0,00 | 3,50 | 29,41 | 0,00 | 29,16 |
|  | Days from peak estrogen (+) |  |  | 8,00 | 48,00 |  | 49,00 | 65,50 |  | 66,50 | 88,25 |  | 8,00 | 80,20 |  |
| Std | Days from peak estrogen (+) |  |  | 0,00 | 25,42 |  | 25,42 | 25,37 |  | 25,37 | 26,85 |  | 0,00 | 29,41 |  |
| Non-birth |  |  |  |  |  |  |  |  |  |  |  |  |  |  |  |
| HH2018 | Days from end-of-cycle (-) |  |  | 114 | 89 |  | 88 | 62 |  | 61 | 43 |  | 114 | 43 |  |
|  | Days from peak estrogen (+) | 120 |  | 8 | 33 | 26 | 34 | 60 | 27 | 61 | 79 | 19 | 8 | 79 | 72 |
|  | Days from end-of-cycle (-) |  |  | 116 | 89 |  | 82 | 62 |  | 61 | 43 |  | 116 | 43 |  |
| HuHu2016 | Days from peak estrogen (+) | 100,38 | 91 | 8 | 41 | 34 | 42 | 62 | 21 | 63 | 81 | 19 | 8 | 81 | 74 |
| HuHu2020 | Days from end-of-cycle (-) |  |  | 88 | 71 |  | 70 | 58 |  | 57 | 43 |  | 88 | 43 |  |
|  | Days from peak estrogen (+) | 92,69 | 86 | 8 | 25 | 18 | 26 | 38 | 13 | 39 | 53 | 15 | 8 | 53 | 46 |
|  | Days from end-of-cycle (-) |  |  | 131 | 97 |  | 96 | 60 |  | 59 | 43 |  | 131 | 43 |  |
| TT2014 | Days from peak estrogen (+) | 108,22 | 103,4 | 8 | 42 | 35 | 43 | 79 | 37 | 80 | 96 | 17 | 8 | 96 | 89 |
| TT2015 | Days from end-of-cycle (-) |  |  | 140 | 83 |  | 82 | 63 |  | 62 | 43 |  | 140 | 43 |  |
|  | Days from peak estrogen (+) | 104,37 | 100,5 | 8 | 65 | 58 | 66 | 85 | 20 | 86 | 105 | 20 | 8 | 105 | 98 |
|  | Days from end-of-cycle (-) |  |  | 137 | 75 |  | 74 | 61 |  | 60 | 43 |  | 137 | 43 |  |
| TT2016 | Days from peak estrogen (+) | 110,14 |  | 8 | 70 | 63 | 71 | 94 | 14 | 85 | 102 | 18 | 8 | 102 | 95 |
| TT2017 | Days from end-of-cycle (-) |  |  | 161 | 83 |  | 82 | 65 |  | 64 | 43 |  | 161 | 43 |  |
|  | Days from peak estrogen (+) | 107,2 | 104,9 | 8 | 86 | 79 | 87 | 104 | 18 | 105 | 126 | 22 | 8 | 126 | 119 |
|  | Days from end-of-cycle (-) |  |  | 145 | 97 |  | 96 | 66 |  | 65 | 43 |  | 145 | 43 |  |
| TT2019 | Days from peak estrogen (+) | 97,47 | 93 | 8 | 56 | 49 | 57 | 87 | 31 | 88 | 110 | 23 | 8 | 110 | 103 |
| Average | Days from end-of-cycle (-) |  |  | 129,00 | 84,75 | 45,25 | 83,75 | 62,13 | 22,63 | 61,13 | 43,00 | 19,13 | 129,00 | 43,00 | 87,00 |
|  | Days from end-of-cycle (-) |  |  | 22,50 | 9,35 | 20,62 | 9,35 | 2,59 | 8,40 | 2,59 | 0,00 | 2,59 | 22,50 | 0,00 | 22,50 |
|  | Days from peak estrogen (+) |  |  | 8,00 | 52,25 |  | 53,25 | 74,88 |  | 75,88 | 94,00 |  | 8,00 | 94,00 |  |
| Std | Days from peak estrogen (+) |  |  | 0,00 | 20,62 |  | 20,62 | 20,51 |  | 20,51 | 22,50 |  | 0,00 | 22,50 |  |
| All cycles |  |  |  |  |  |  |  |  |  |  |  |  |  |  |  |
| Average | Days from end-of-cycle (-) |  |  | 124,06 | 83,65 | 41,35 | 82,65 | 62,59 | 21,06 | 61,59 | 43,00 | 19,59 | 121,89 | 43,00 | 79,84 |
|  | Days from end-of-cycle (-) |  |  | 21,40 | 7,36 | 19,38 | 7,36 | 3,24 | 6,30 | 3,24 | 0,00 | 3,24 | 22,27 | 0,00 | 22,21 |
|  | Days from peak estrogen (+) |  |  | 8,00 | 48,41 |  | 49,41 | 69,47 |  | 70,47 | 89,06 |  | 8,00 | 86,89 |  |
| Std | Days from peak estrogen (+) |  |  | 0,00 | 19,42 |  | 19,42 | 20,01 |  | 20,01 | 21,40 |  | 0,00 | 22,27 |  |

16 Supplementary table 6: additional data on the lenght of the different CLD phases per giant panda.

| Cycle | Category | Latitude | Average anestrus bodyweight | Average postestrus bodyweight | A body weight | Dormancy I |  |  | Dormancy II |  |  | Dormancy III |  |  | Total CL dormancy |  |  |  |  |  |
| --- | --- | --- | --- | --- | --- | --- | --- | --- | --- | --- | --- | --- | --- | --- | --- | --- | --- | --- | --- | --- |
|  |  |  |  |  |  | Start | Stop | Interval | Start | Stop | Interval | Start | Stop | Interval | Start | Stop | Interval |  |  |  |
| 58569 | 2014 NB | Days from end-of cycle (-) |  |  |  |  | 131 | 97 |  | 96 | 60 |  | 59 | 43 |  | 131 | 43 |  |  |  |
|  |  | Days from peak estrogen (+) |  |  |  |  | 8 | 42 | 35 | 43 | 79 | 37 | 80 | 96 | 17 | 8 | 96 | 89 |  |  |
|  |  | Days from end-of cycle (+) |  | 108,22 | 103,40 | 4,82 |  | 140 | 83 |  | 82 | 63 |  | 82 | 43 |  | 140 | 43 |  |  |
|  | 2015 NB | Days from end-of cycle (-) |  |  |  |  | 8 | 65 | 58 | 66 | 85 | 20 | 86 | 105 | 20 | 8 | 105 | 98 |  |  |
|  |  | Days from peak estragen (+) |  | 104,37 | 100,50 | 3,87 |  | 137 | 75 |  | 74 | 61 |  | 60 | 43 |  | 137 | 43 |  |  |
|  |  | Days from end-of cycle (+) |  |  |  |  | 8 | 70 | 63 |  | 71 | 84 | 14 | 85 | 102 | 18 | 8 | 102 | 95 |  |
|  | 2016 NB | Days from end-of cycle (-) |  |  |  |  | 161 | 83 |  | 82 | 65 |  | 64 | 43 |  | 161 | 43 |  |  |  |
|  |  | Days from peak estrogen (+) |  | 110,14 |  |  | 8 | 63 |  | 64 | 42 |  | 63 | 100 |  | 8 | 100 |  |  |  |
|  |  | Days from end-of cycle (+) |  | 107,20 | 104,90 | 2,30 |  | 8 | 86 | 79 |  | 87 | 104 | 18 | 105 | 126 | 22 | 8 | 126 | 119 |
|  | 2018 PP | Days from end-of cycle (-) |  |  |  |  | 144 | 89 |  | 88 | 70 |  | 69 | 43 |  | 144 | 43 |  |  |  |
|  |  | Days from peak estrogen (+) |  | 99,95 | 94,50 | 5,45 |  | 8 | 63 | 55 | 64 | 42 | 19 | 63 | 100 | 27 | 8 | 100 | 101 |  |
| Days from end-of cycle (+) |  |  |  |  |  | 145 | 97 |  | 96 | 66 |  | 65 | 43 |  | 145 | 43 |  |  |  |  |
| 58723 | 2019 NB | Days from peak estrogen (+) |  | 97,47 | 93,00 | 4,47 |  | 8 | 56 | 49 |  | 57 | 87 | 31 | 88 | 110 | 23 | 8 | 110 | 103 |
|  |  | Days from end-of cycle (-) |  | 104,56 | 99,26 | 4,18 |  | 143,00 | 87,33 | 56,50 | 86,33 | 64,17 | 23,17 | 63,17 | 43,00 | 21,17 | 143,00 | 43,00 | 100,83 |  |
|  |  | Days from end-of cycle (+) |  | 4,96 | 5,30 | 1,20 |  | 10,18 | 8,71 | 14,64 | 8,71 | 3,66 | 8,84 | 3,66 | 0,00 | 10,18 | 0,00 | 10,17 |  |  |
|  | Average | Days from peak estrogen (+) |  |  |  |  | 8,00 | 63,67 | 64,67 | 86,83 |  |  | 87,83 | 108,00 |  | 8,00 | 108,00 |  |  |  |
|  |  | Std |  |  |  |  | 0,00 | 14,62 |  | 14,62 | 8,84 |  | 8,84 | 10,18 |  | 0,00 | 10,18 |  |  |  |
|  |  | 58725 | Days from end-of cycle (-) |  |  |  |  | 116 | 83 |  | 82 | 62 |  | 61 | 43 |  | 116 | 43 |  |  |
|  | Days from peak estrogen (+) |  |  | 100,38 | 91,00 | 9,38 |  | 8 | 41 | 34 |  | 42 | 62 | 21 | 63 | 81 | 19 | 8 | 81 | 74 |
|  | Days from end-of cycle (+) |  |  |  |  |  | 127 | 84 |  | 83 | 60 |  | 59 | 43 |  | 127 | 43 |  |  |  |
|  | 2020 NB | Days from peak estrogen (+) |  | 94,82 | 93,50 | 1,32 |  | 8 | 51 | 44 |  | 52 | 75 | 24 | 76 | 92 | 17 | 8 | 92 | 85 |
|  |  | Days from end-of cycle (-) |  |  |  |  | 88 | 71 |  | 70 | 58 |  | 57 | 43 |  | 88 | 43 |  |  |  |
|  |  | Days from peak estrogen (+) |  | 92,69 | 86,00 | 6,69 |  | 8 | 25 | 18 | 26 | 38 |  | 27 | 33 | 15 | 8 | 33 |  |  |
| 58741 | 2015 PS | Days from end-of cycle (-) |  | 95,96 | 90,17 | 5,80 |  | 110,33 | 79,33 | 39,00 | 78,33 | 60,00 | 22,50 | 59,00 | 43,00 | 18,00 | 110,33 | 43,00 | 79,50 |  |
|  |  | Days from peak estragen (+) |  | 3,97 | 3,82 | 4,10 |  | 20,11 | 7,23 | 7,07 | 7,23 | 2,00 | 2,12 | 2,00 | 0,00 | 1,41 | 20,11 | 0,00 | 7,78 |  |
|  |  | Std |  |  |  |  | 8,00 | 39,00 |  | 40,00 | 58,33 |  | 59,33 | 75,33 |  | 8,00 | 75,33 |  |  |  |
|  | Average | Days from peak estrogen (+) |  |  |  |  | 0,00 | 13,11 |  | 13,11 | 18,77 |  | 18,77 | 20,11 |  | 0,00 | 20,11 |  |  |  |
|  |  | Std |  |  |  |  |  |  |  |  |  |  |  |  |  |  |  |  |  |  |
|  |  | 58743 | Days from end-of cycle (-) |  |  |  |  |  |  |  |  |  |  |  |  |  |  | 124 | 43 |  |
|  | Days from peak estrogen (+) |  |  |  |  |  |  |  |  |  |  |  |  |  |  |  | 8 | 89 | 82 |  |
|  | Days from end-of cycle (+) |  |  | 120,00 |  |  |  | 106 | 80 |  | 79 | 61 |  | 60 | 43 |  | 106 | 43 |  |  |
|  | 2016 NB | Days from peak estrogen (+) |  | 120,00 |  |  |  | 8 | 34 | 27 | 35 | 53 | 19 | 54 | 71 | 18 | 8 | 71 | 64 |  |
|  |  | Days from end-of cycle (-) |  |  |  |  | 114 | 89 |  | 88 | 62 |  | 61 | 43 |  | 114 | 43 |  |  |  |
|  |  | Days from peak estrogen (+) |  | 120,00 |  |  | 8 | 33 | 26 | 34 | 60 | 27 | 61 | 79 | 19 | 8 | 79 | 72 |  |  |
| 2019 P | Days from end-of cycle (-) |  |  |  |  | 114 | 74 |  | 73 | 57 |  | 56 | 43 |  | 114 | 43 |  |  |  |  |
|  | Days from peak estrogen (+) |  | 120,00 |  |  | 8 | 48 | 41 | 49 | 65 | 17 | 66 | 79 | 14 | 8 | 79 | 72 |  |  |  |
|  | Days from end-of cycle (+) |  | 120,00 |  |  |  | 111,33 | 81,00 | 31,33 | 80,00 | 60,00 | 21,00 | 59,00 | 43,00 | 17,00 | 111,33 | 43,00 | 69,33 |  |  |
| 58884 | 2017 PS | Std |  |  |  |  | 4,62 | 7,55 | 8,39 | 7,55 | 2,65 | 5,29 | 2,65 | 0,00 | 2,65 | 4,62 | 0,00 | 4,62 |  |  |
|  |  | Days from peak estrogen (+) |  |  |  |  | 8,00 | 38,33 |  | 39,33 | 58,33 |  | 60,33 | 76,33 |  | 8,00 | 76,33 |  |  |  |
|  |  | Std |  |  |  |  | 0,00 | 8,39 |  | 8,39 | 6,03 |  | 6,03 | 4,62 |  | 0,00 | 4,62 |  |  |  |
|  | 2018 PS | Days from end-of cycle (-) |  |  |  |  |  |  |  |  |  |  |  |  |  |  | 83 | 43 |  |  |
|  |  | Days from peak estrogen (+) |  |  |  |  |  |  |  |  |  |  |  |  |  |  | 8 | 48 | 41 |  |
|  |  | Days from end-of cycle (+) |  | 106,02 | 97,14 | 8,88 |  | 94 | 77 |  | 76 | 62 |  | 61 | 43 |  | 94 | 43 |  |  |
|  | 2019 PS | Days from peak estrogen (+) |  | 108,58 | 104,45 | 4,13 |  | 8 | 25 | 18 | 26 | 40 | 15 | 41 | 59 | 19 | 8 | 59 | 52 |  |
|  |  | Days from end-of cycle (-) |  |  |  |  | 107 | 87 |  | 86 | 67 |  | 66 | 43 |  | 107 | 43 |  |  |  |
|  |  | Days from peak estrogen (+) |  | 108,39 | 103,30 | 5,09 |  | 8 | 28 | 21 | 29 | 48 | 20 | 49 | 72 | 24 | 8 | 72 | 65 |  |
|  | 2020 P | Days from end-of cycle (-) |  |  |  |  | 97 | 83 |  | 82 | 62 |  | 61 | 43 |  | 97 | 43 |  |  |  |
|  |  | Days from peak estrogen (+) |  | 109,56 | 102,09 | 7,47 |  | 8 | 22 | 15 | 23 | 43 | 21 | 44 | 62 | 19 | 8 | 62 | 55 |  |
| Days from end-of cycle (+) |  |  | 108,14 | 101,75 | 6,39 |  | 99,33 | 82,33 | 18,00 | 81,33 | 63,67 | 18,67 | 62,67 | 43,00 | 20,67 | 95,25 | 43,00 | 53,25 |  |  |
| 58888 | 2019 PS | Std |  |  |  |  | 6,81 | 5,03 | 3,00 | 5,03 | 2,89 | 3,21 | 2,89 | 0,00 | 2,89 | 9,88 | 0,00 | 9,88 |  |  |
|  |  | Days from peak estrogen (+) |  |  |  |  | 8,00 | 25,00 |  | 26,00 | 43,67 |  | 44,67 | 64,33 |  | 8,00 | 60,25 |  |  |  |
|  |  | Std |  |  |  |  | 0,00 | 3,00 |  | 3,00 | 4,04 |  | 4,04 | 6,81 |  | 0,00 | 9,88 |  |  |  |
|  | 2019 P | From end-of cycle |  |  |  |  | 140 | 90 |  | 89 | 64 |  | 63 | 43 |  | 140 | 43 |  |  |  |
|  |  | From peak estrogen |  | 81,28 | 81,00 | 0,28 | 8 | 58 | 51 | 59 | 84 | 26 | 85 | 105 | 21 | 8 | 105 | 98 |  |  |
|  |  | Days from end-of cycle (+) |  |  |  |  |  |  |  |  |  |  |  |  |  |  |  |  |  |  |
|  | 58941 | Days from end-of cycle |  |  |  |  | 148 | 80 |  | 79 | 64 |  | 63 | 43 |  | 148 | 43 |  |  |  |
|  |  | From peak estragen |  |  |  |  | 8 | 76 | 69 | 77 | 92 | 16 | 93 | 113 | 21 | 8 | 113 | 106 |  |  |
|  |  | Days from end-of cycle (+) |  | 109,50 | 106,33 | 3,17 |  |  |  |  |  |  |  |  |  |  |  |  |  |  |

17
18
19

20
21

Supplementary table 7: information on husbandry and diet for each female giant panda.

| Studbooknr | SB569 | SB723 | SB741 | SB884 | SB868 | SB941 |
| --- | --- | --- | --- | --- | --- | --- |
| Zoological Institution | Edinburgh Zoo - RZSS | Zooparc de Beauval | Pairi Daiza | Ouwehand | Berlin Zoo | Ähtäri Zoo |
| Indoor temperature (°C) | 8-18°C | 8-20°C | 8-20°C | 15-22°C | 10-22°C | 8-22°C |
| Outdoor Enclosure (m<sup>2</sup>) | 1500 | n.a. | 681 | 1200 | 1130 | 1600 |
| Indoor Enclosure (m<sup>2</sup>) | 80 | n.a. | 121 | 300 | 80 | 199 |
| Backstage Enclosure (m<sup>2</sup>) | 40 | n.a. | 34 | 25 | 52 | 27 |
| Indoor humidity (%) | n.a. | n.a. | n.a. | > 70% | 50% | 55-57% |
| Bamboo offered (kg/day) | 40-90 | n.a. | n.a. | 40-45 | 33 | 50 |
| Bamboo consumed (kg/day) | 10-50 | 14 | n.a. | n.a. | 9 | n.a. |
| Predominant bamboo species | *P. Aurata*, *P. bissettii* | *P. Bissetti*, *P. Humilis*, *P. Viridiglaucescens*, *P. Nuda*, *P. Flexuosa*, *P. Aureocaulis*, *P. Aureosulcata*, *P. Japonica*, *P. Vicax*, *P. Nigra* | *P. Japonica*, *P. Bissetti*, *P. Flexuosa*, *P. Spectabilis*, *P. Fastuosa* | *P. Bissetti*, *P. Fastuosa*, *P. Japonica*, *P. Viridi G*, *P. Vivax*, *P. Shaghai*, *P. Aureosulcata*, *P. Spectabilis*, *P. Dacora*, *P. Humilis*, *P. Palmata*, *P. Boreana*, *P. Henonis*, *P. Iridescentis*, *P. Nigra*, *P. Aurea* | *P. japonica*, *P. vivax*, *'Spectabilis'*, *P. vivax*, *P. aureosulcata*, *'Spectabilis'*, *P. iridescentis*, *P. vivax*, *'Shanghai 3'*, *P. bissettii*, *P. viridiglaucescens*, *P. aureosulcata*, *'Aureocaulis'*, *P. humilis*, and *P. aurea*, *P. viridis*, *Bashania fargesii*, *P. nigra* | *P. Bissetti*, *B. Humilis*, *P. Japonica*, *P. Aureosulcata*, *spectabilis* |
| Cake (C) /Pellets (P) (g/day) | 2013-2019: 500 (P), 2019-....: 500 (C) | 600-700 (P) | 800 (C) | 1000 (C) | 700 (C) | 750 (C) |
| Apples (g/day) | 150-300 (training)<br>50 - 100 (training) | 150 (training) | 250<br>500 | 100<br>600 | 150-250<br>100-300 (incl. beetroots, sweet potato) | 140<br>500 |
| Carrots (g/day) |  |  |  |  |  |  |
| Supplements | Vit D, calcium | n.a. | Vit D, calcium | Vitamins, Calcium |  | Vitamins |

22
23
24
